# Host adaptation and genome evolution of the broad host range fungal rust pathogen, *Austropuccinia psidii*

**DOI:** 10.1101/2025.05.29.656941

**Authors:** Zhenyan Luo, Peri Tobias, Lavi Singh, Chongmei Dong, Alyssia M Martino, Maria Quecine, Nelson Massola, Lilian Amorim, Jianbo Li, Smriti Singh, Ziyan Zhang, Ashley Jones, Robert F Park, Benjamin Schwessinger, Richard Edwards, Thais Boufleur

## Abstract

Rust diseases on plants are caused by fungi in the order Pucciniales. Typically, rust fungi have narrow host specificity however the pandemic biotype of *Austropuccinia psidii* has an unusually broad host range causing disease on over 480 myrtaceous species globally. We assembled and analysed a fully phased chromosome-level genome for the pandemic *A. psidii* and addressed key outstanding questions of its infection biology. Our research revealed a conserved rust fungal karyotype of 18 haploid chromosomes, in line with fungi for distantly related cereal rusts. We observed chromosomal re-assortment between the two nuclei, with one nucleus carrying 19 and the other 17 chromosomes. The synteny of universal single-copy orthologs is mostly maintained with the distantly related rust fungus *Puccinia graminis* f. sp. *tritici*. In contrast, nucleotide composition and methylation profiles of *A. psidii* are distinct compared to rust fungi with smaller genome sizes that have not undergone massive transposable element expansions. Our analysis of mating type loci supports a tetrapolar mating system for *A. psidii* with a novel finding of expanded numbers of pheromone peptide precursors. We show that infection dynamics of *A. psidii* are consistent on four different susceptible host species separated by 65 mya of evolution and that transcriptional regulation during infection reveals two distinct waves of gene expression in early and late infection, including allele-specific expression of candidate effectors. Together, these findings enhance the understanding of the genome biology and pathology of *A. psidii*, while also providing a valuable resource for future research on this serious rust pathogen.

## 1. Introduction

Rust diseases, caused by obligate biotrophic fungi from the order Pucciniales, have historically devastated economies and contributed to famines worldwide (1). The *Pucciniales* are one of the largest plant pathogenic fungal orders (2), and although they typically display a high degree of host specificity due to coevolution (3) , species such as *Phakopsora pachyrhizi*, *P. meibomiae*, and *Austropuccinia psidii* infect a wide range of hosts within leguminous plants (4–6) and the Myrtaceae (7), respectively.

Originally classified as a monospecific genus, *Austropuccinia* was defined by a single species *A. psidii* (8) until 2024 when *A. licaneae* (Henn) Ebinghaus & Dianese*, comb. nov.*, was taxonomically proposed as a fungus that infects *Licania* trees from the Chrysobalanaceae family in the Amazons (9). The broad host range of *A. psidii* is not universally applicable to all pathogen isolates and biotype. Within its native range of South America most biotypes exhibit various degrees of host specificity, for example the guava biotype does not sporulate on *Syzygium jambos* (10–13). Outside its native range the behaviour of biotypes might be different. For example, a recent study showed that the South American eucalyptus biotype of *A. psidii* produced infection symptoms on several New Zealand myrtaceous species not found in South America (14). Further, the pandemic biotype of *A. psidii* has been shown to infect over 480 Myrtaceae globally (11, 15), and, as a highly invasive pathogen, has caused dramatic impacts to natural ecosystems notably in Australia, New Zealand, and Hawaii (16–18). *Austropuccinia psidii*, like other rust fungi, exists predominantly in the asexual dikaryotic urediniospore stage of its life cycle, with two separate haploid nuclei coexisting in the same cytoplasm. A key goal in eukaryotic genome evolution research is achieving fully phased genome assemblies with telomere to telomere (T2T) chromosomes. Rust fungi genome sizes ranges from ∼70 Mb to ∼3 Gb (19, 20) and long reads alone are insufficient to assemble and phase dikaryotic genomes completely especially of sizes greater 1 Gb (21). Until very recently, the most effective approach combines PacBio High-Fidelity (HiFi) reads (22) with Oxford Nanopore Technologies (ONT) ultra-long reads, and long-range chromatin interaction data such as Hi-C (Illumina) (23). Currently, the few rust fungal genomes that have been fully phased and assembled to near-T2T are primarily host-specific cereal rusts with a karyotype of 18 chromosomes (24–30). A partially phased genome for the pandemic lineage of *A. psidii* based on PacBio RSII and Sequel technologies revealed over 90 % transposable elements (TEs), unusually large telomeres, and a haploid genome size of around 1 Gb. However, scaffolding and phasing of this *A. psidii* genome (v1) (21) was challenging due to early-generation PacBio read error and long read assembly software. The v1 genome therefore incorporated a considerable number of residual errors, especially around highly repeat-rich regions like centromeres and mating-type loci.

The large genome size of *A. psidii* is hypothesized to result from bursts of TEs and their potential silencing via methylation and cytosine deamination which might have led to the comparatively low GC content, at 33.8 % (21). Understanding the mechanisms behind genome size expansion in fungal pathogens and how TE activity contributes to this phenomenon, are critical questions in fungal biology and evolution (31). TEs are repeated sequences that enable their own proliferation within the genome (31), with roles in chromosome rearrangement and gene expression regulation (32). Because uncontrolled proliferation can lead to genome instability (33), fungi have evolved defence mechanisms that include RNA interference (RNAi), repeat-induced point mutation (RIP), DNA or histone methylation (34–37). How these TE silencing mechanisms contribute to genome evolution in rust fungi including *A. psidii* is currently poorly understood.

TEs also play an important role in mating-type loci evolution in rust fungi (38). Genes at the mating-type (MAT) loci regulate mating in fungi (39), which exhibit either bipolar or tetrapolar mating systems (40). In most Basidiomycota, successful mating requires the presence of heterozygous pheromone and receptor (PR) and homeodomain (HD) loci (41). In rust fungi, the PR locus consists of one pheromone gene (*Pra*) and at least one pheromone peptide precursor gene (*mfa*), while the HD locus contains two tightly linked homeodomain transcription factor genes (*bW-HD1* and *bW-HD2*) (40, 42). Using an early version of the near-T2T *A. psidii* genome (43), Ferrarezi et al. (2022) were able to identify the *Pra* and *HD* genes but no putative *mfa* genes. Further analyses revealed that this species most likely has a tetrapolar mating system characterized by a multiallelic HD locus and a biallelic PR locus, facilitating outcrossing on universally susceptible hosts (38, 44). The MAT loci were poorly resolved in the v1 *A. psidii* genome, meaning that detailed information available on the genome biology and linkage of mating-type loci in *A. psidii* was lacking. Recent studies on phased genomes of cereal rust species supported the hypothesis of a tetrapolar mating system in rust fungi by revealing the MAT loci on two different chromosomes, multiallelic HD genes and a biallelic PR locus, with the exception of *P. graminis* f. sp. *tritici*, which potentially possesses multiple alleles at the PR locus (38). The pandemic lineage of *A. psidii* has an expanding list of host species partly explained by its global spread (7, 15, 45, 46), and the results of several studies suggest that it has different infection dynamics on different hosts (47–50). While symptoms across multiple host species in controlled inoculations are reported to vary considerably (14, 50, 51), to our knowledge no study has investigated infection dynamics only within highly susceptible host plants of different species. We investigated this by testing infection dynamics and pustule development within previously determined susceptible individuals from four species separated by 65 mya of evolution.

This research confirms the conserved karyotype in rust fungi, identifies the unusual chromosome assortment and addresses four fundamental knowledge gaps around *A. psidii* biology that are important to develop long-term pathogen containment. Based on these existing knowledge gaps combined with new resources for *A. psidii*, we posed the following six key questions to investigate, 1) does *A. psidii* have distinct infection progression in different susceptible host species? 2) what is the karyotype of *A. psidii*? 3) what is the conserved gene synteny compared to distantly related rust fungi? 4) does DNA methylation play a role in TE silencing? 5) what is the genome biology of mating-type loci? and 6) how prevalent allele specific expression during the infection process?

## 2. Materials and Methods

### 2.1 Infection assay

Plants from four different species belonging to the Myrtaceae family were used in this experiment, including *Leptospermum scoparium*, *Melaleuca alternifolia*, *Melaleuca quinquenervia,* and *Syzygium jambos*. The plants were selected based on confirmed susceptible phenotype from controlled inoculations (50, 52). Plants were cultivated in a peat and perlite mixture (1:5) and grown in a greenhouse at controlled temperatures of 24 °C during the day and 20 °C at night on a 12 h cycle.

*Austropuccinia psidii* urediniospores of isolate Au_3 (21), which belongs to the pandemic biotype, were amplified on *S. jambos* and used for all subsequent infection experiments. Urediniospores from five-days-old pustules were harvested by shaking, suspended in 0.05 % Tween 20, and adjusted to 10^6^ urediniospores ml^-1^. The urediniospore suspension was painted on young leaves of the second internode (both abaxial and adaxial surfaces) for all plants, as described previously (47). Plants were incubated for 24 h in a dark sealed box with plentiful water to maintain high humidity, at 23 °C, and subsequently moved to a growth chamber and maintained at 22 ± 2 °C, 60 % relative humidity, and a 14 h light photoperiod. Three biological replicates were inoculated for each species, with each biological replicate corresponding to a distinct plant individual.

To compare infection symptom development, multiple selected leaves were manually inspected from one to 12 days post-inoculation (dpi) to register the occurrence of the first visible symptoms, along with recording images of all leaves. Both adaxial and abaxial surfaces of individual leaves were documented at a distance of 1 - 2 cm.

### 2.2 Cytological study

Fresh urediniospores were suspended in sterilized ddH2O containing 0.05 % Tween 20 at a concentration of 1 mg/mL and mixed by inverting gently. The suspension was spread into a thin layer on 2 % water agar plates. The plates were covered with foil and incubated for 24 h at 18 °C to stimulate germination. Germinated urediniospores were suspended in ddH2O and centrifuged at 5,000 rpm for 2 min to remove ddH2O. Following treatment with nitrous oxide gas at 1.0 MPa for 1.5 h, the urediniospores were fixed in ice-cold 90 % acetic acid for 8 min and then centrifuged at 5,000 rpm for 2 min to remove the acetic acid. The tubes were placed on ice and washed twice with distilled water and then centrifuged at 5,000 rpm for 2 min to remove residual water. The urediniospores were digested in 1.5 % Lysing Enzymes from *Trichoderma harzianum* (Cat. No. L1412, Sigma) for 2 h at 28 °C. After digestion, they were washed twice with 70 % ethanol and then centrifuged at 5,000 rpm for 2 min to remove the enzymes and ethanol. The urediniospores were squashed using a fine needle in 50 µL of 70 % ethanol, centrifuged at 5,000 rpm for 2 min, and dried to remove the remaining ethanol. The urediniospores were vortexed at maximum speed in 50 µL of glacial acetic acid to separate cells from one another. The cell suspension was dropped on microscope slides in a humid box and dried slowly.

Chromosomes were observed with a Zeiss Axio Imager epifluorescence microscope, and images were captured with a Retiga EXi CCD camera (QImaging, Surrey, BC, Canada) operated with Image-Pro Plus version 7.0 software (Media Cybernetics Inc., Bethesda, MD, USA).

### 2.3 PacBio and Oxford Nanopore Technologies (ONT) long-read DNA sequencing

To improve the scaffolding on the previous assembled *A. psidii* (Au3_v1) genome (21), 16 µg of previously obtained High Molecular Weight (HMW) DNA was sent to the Australian Genome Research Facility (AGRF) at the University of Queensland, Brisbane, Australia, for PacBio Sequel II HiFi sequencing. For ONT sequencing, HMW DNA of Au3 urediniospores preserved at -80 °C was extracted using a modified Cetrimonium Bromide (CTAB) extraction protocol (21, 53). The extracted DNA was sent to the Biomolecular Resource Facility (BRF) at the Australian National University (ANU). The DNA was size selected using a BluePippin (Sage Science) with a cutoff of 40kb+ for library preparation according to the manufacturer’s protocol, with the Ligation Sequencing kit V14 (SQK-LSK114). Sequencing was performed on a PromethION using a flow cell FLO-PRO114M according to the manufacturer’s instructions. The pbccs v.6.4.0 was used to generate HiFi reads with tag ‘—mini-rq=0.88’. Actc v0.6.0 (https://github.com/PacificBiosciences/actc) was applied to map subreads to HiFi reads, followed by processing with DeepConsensus v.1.2.0 for read corrections (54). PromethION fast5 reads were basecalled to fastq with Guppy v.6.4.6 (https://community.nanoporetech.com) with model dna_r10.4.1_e8.2_400bps_hac@v3.5.2. Sequencing quality was checked with Nanoplot v. 1.41.0 (55). ONT reads were processed with Chopper v.0.5.0 (55) retaining the longest 35 x fraction and reads with a quality score of ≥ Q7.

### 2.4 Illumina and ONT RNA sequencing

For Illumina RNA sequencing, *S. jambos* plants with young growing tissues were inoculated with *A. psidii* Au3 isolate as described in 2.1. Six fragments of infected leaves located at the first to third internode were collected at one, two, three, four, five and six dpi, snap frozen in liquid nitrogen and stored at -80 °C for total RNA extraction. *Leptospermum scoparium*, *M*. *alternifolia*, *M. quinquenervia* and *S. luehmannii* were sampled following the similar procedure at three, six, and nine dpi. Whole leaves were collected for these species due to their smaller leaf sizes.

Total RNA was extracted from triplicate samples of plant-infected leaf tissues and ungerminated urediniospores of the Au3 isolate. The samples were homogenized using TissueLyser II (25 Hz for 2 mins) and the RNA was extracted with the Spectrum^TM^ plant total RNA kit (Sigma) as per manufacturer’s instructions. DNase treatment was carried out using TURBO DNA-freeTM Kit (Thermo Fisher Scientific). Oligo d(T) 25 magnetic beads (New England Biolabs) were used to purify the RNA as per manufacturer instructions. The RNA quantity was assessed using a Qubit RNA kit on a Qubit Fluorometer (Thermo Fisher Scientific). The quality of RNA was determined using a Nanodrop (Molecular Devices) and the integrity was checked by Gel electrophoresis (1 % agarose gel). Purified RNA was sent to Azenta Life Sciences for RNA-sequencing with strand-specific polyA mRNA selection using 150 bp paired-end Illumina NovaSeq and a sequencing depth of 20 million reads.

For ONT direct RNA sequencing, 3 mg spores per 1 mL Novec^TM^ 7100 engineered fluid (3MTM) was sprayed onto plants of *S. jambos* with more than 6 juvenile, pink leaves. Plants were then incubated as described in 2.1. At three and six dpi, 2 to 3 grams per infected leaves were collected, snap frozen in liquid nitrogen and stored at -80 °C. Total RNA was extracted using an adapted CTAB protocol (56, 57), resuspended in 10 mM Tris-HCl pH 7.5 and stored at -80 °C. A NanoDrop spectrophotometer and Qubit RNA Broad Range Assay Kit (both by Thermo Fisher Scientific) were used for RNA quantification and quality control, and the integrity was checked by Gel electrophoresis (1 % agarose gel). Dynabeads Oligo (dT)25 (Invitrogen) was used to purify 2 µg poly(A) tailed RNA (mostly mRNA) from total RNA following the manufacturer’s instructions. Quality control was performed as described above. Approximately 500 ng of poly(A) transcripts were used as input for the direct RNA sequencing library. RNA reverse transcription was done with Induro Reverse Transcriptase following the Induro Reverse Transcriptase Protocol (NEB M0681). The remaining library preparation and ONT Direct RNA sequencing kit (SQK-RNA002) were performed using FLO-MIN106D MinION flow cell as per ONT manufacturer’s instructions. Approximately 2 million reads per library were obtained. The raw RNA sequencing signals captured in ONT fast5 files were base called with Guppy v.6.4.2 (Oxford Nanopore Technologies) using the high accuracy config model (rna_r9.4.1_70bps).

### 2.5 Genome assembly and scaffolding with Hi-C data

The data were assembled using 18.5 X of HiFi and 39.5 X of Nanopore read depths per haplotype, supplemented with 80 Gb of raw Hi-C data (43), utilizing Hifiasm software (v.0.19.5-r592) (58, 59) using default settings. HiFi and ONT reads were mapped individually to the draft assembly with MiniMap2 v2.24 (60). The per-contig coverage was estimated with BBMap v.38.90 pileup.sh script (https://github.com/BioInfoTools/BBMap/tree/master). Contigs with low coverage were extracted with SAMtools v.1.1 (61), and HiFi and ONT reads that mapped to these contigs were identified and removed from the fastq files with BBmap v.38.93 (https://github.com/BioInfoTools/BBMap/tree/master). A new assembly was run with the filtered reads as described above.

The two-phased genome outputs were merged, and scaffolding was performed using Hi-C data, following the Aiden Lab pipelines (https://github.com/aidenlab). The merged file was run through the Juicer pipeline (v.1.6) with default parameters (62, 63). The final output from Juicer was used with the 3D-DNA pipeline (v.180922) with the following parameters ‘-m diploid –build-gapped-map --sort-output’ (62, 63). The output of 3D-DNA was manually curated within the Juicebox visualization software (v.1.11.8 for Mac) (62, 63), and the revised assembly file was resubmitted to the 3D-DNA post review pipeline with the following parameters ’—build-gapped-map –sort-output’ for final assembly and FASTA files. For post-scaffolding contamination removal and clean-up, unplaced contigs were BLASTed against the RefSeq database of NCBI for contaminant identification.

Functional completeness of each haploid genome and from the dikaryon was estimated using BUSCO v.5.4.0 (64) with the basidiomycota_odb10 lineage database. The most complete haplotype was named hapA and the chromosomes were arranged by size and each starting with the short chromosome arm. HapB was mapped to hapA to identify and name homologous chromosomes and organize unscaffolded contigs by size with PAFScaff v.0.6.3 (65). Terminal telomeric repeats were predicted using Telociraptor v0.8.0 (https://github.com/slimsuite/telociraptor). The final assembly was designated as the v3 assembly (hapA, hapB and mt).

### 2.6 Repeat and gene annotation

RepeatMasker v.4.0.9 was applied to reannotate the genome with a *de novo* constructed TE database from Au3_v1 (21). The Wicker classification system (65) was applied to classify TEs into the superfamily level. For TE families that could not be classified by REPET v.3, new classifications were assigned based on TE sequences showing at least 70 % identity to the consensus sequences of these TE families. For calculating TE density, all insertions were included in the analysis. To avoid bias in analysing the relationship between TEs and methylation, only insertions longer than 1kbp were included.

The haploid genomes were soft-masked and functionally annotated using the Funannotate v.1.8.15 pipeline (https://zenodo.org/record/2604804). The prediction includes “train”, “predict” and “update”. The transcriptome generated in 2.6 was used in the “train” step, and ‘--jaccard_clip’ was added for better fungal annotation. In the “predict” step, UniProt database was added as protein hints for the first round, the weight of each gene prediction tools was set as the default. In the “update” step, the parameter of ‘--jaccard_clip’ was added. After the first round of annotation, protein files generated in opposite haplotypes were added as protein hints in the “predict” step for the second round.

Given the repeat-rich nature of the *A. psidii* genome the predicted genes might be related to TEs despite soft-masking, which could influence subsequent analysis. To address this, Orthofinder v.2.5.5 (66) was employed to identify orthogroups between *A. psidii*, *Pst 104E* and *Pt 76*. Most orthogroups have less than eight gene members and less than 20 orthogroups of *Pst 104E* and *Pt 76* have more than 20 gene members, so orthogroups containing more than 20 gene members in *A. psidii* were classified as repeat-related genes and were removed from further analysis.

The predicted genes were functionally annotated using the ‘funannotate annotate’ command in the pipeline and described as follows. Protein-coding gene models were used to pull annotations from Pfam, dbCAN (CAZyme), and Gene Ontology (GO) databases. GO and Pfam enrichment analysis were implemented using ‘enricher’ function in clusterProfiler v.4.12.6 R package (67), which fits a hypergeometric test to identify over-represented terms (Benjamini-Hochberg FDR of p < 0.05). To predict the secretome of the *A. psidii* isolate Au3, proteins containing a signal peptide cleavage site were identified with SignalP v.6.0 (68). Subsequently, proteins with transmembrane (TM) domains and glycosylphosphatidylinositol (GPI) anchoring were identified with DeepTMHMM v.1.0.39 (69) and NetGPI v.1.1 (70). Proteins with a signal peptide cleavage site and absence of TM-domain and GPI-anchors were predicted to be part of the *A. psidii* secretome. The subcellular localization of the secretome was predicted using WoLF PSORT (71). Finally, Blastp v.2.12.0+ (72, 73) was employed to detect secreted proteins that were either shared among haplotypes or hemizygous, applying criteria of at least 90 % identity and 50 % query coverage. Blast results were checked manually and the best hit in both haplotypes was used as a criterion to identify the alleles. Secreted proteins that were differentially expressed in planta when compared to ungerminated urediniospores identified in 2.11, were classified as the candidate effectors (CEs) of *A. psidii*. The pattern of expression of the identified alleles (shared) and specific (hemizygous) CEs was plotted with pheatmap R package v.1.0.12 (74).

### 2.7 Chromosome synteny analysis

Chromosome synteny analysis was performed using single-copy “Complete” genes from the BUSCO genome completeness runs. For this analysis, one copy of Chr14 was placed in hapB. Synteny blocks between pairs of assemblies were determined by collinear runs of matching BUSCO gene identifications. For each assembly pair, BUSCO genes rated as “Complete” in both genomes were ordered and oriented along each chromosome. Synteny blocks were then established as sets of BUSCO genes that were collinear and uninterrupted (i.e., sharing the same order and strand) in both genomes, starting at the beginning of the first BUSCO gene and extending to the end of the last BUSCO gene in the block. We note that local rearrangements and breakdowns of synteny between BUSCO genes will not be identified and may be falsely marked as syntenic within a synteny block.

Chromosome synteny was then visualized by arranging chromosomes for each assembly in rows and plotting the synteny blocks between adjacent assemblies. In each case, the chromosome order and orientation were set for one “focus” assembly, and the remaining assemblies arranged to maximize the clarity of the synteny plot, propagating out from the focal genome. For each chromosome, the best hit in its adjacent assembly was established as that with the maximum total length of shared synteny blocks, and an anchor point established as the mean position along the best hit chromosome of those synteny blocks. Where the majority of synteny was on the opposite strand, the chromosome was reversed and given an “R” suffix in the plot. Chromosomes were then ordered according to their best hits and, within best hits, the anchor points. Synteny blocks sharing the same orientation were plotted blue, whilst inversions are plotted in red. Code for establishing and visualizing chromosome synteny based on shared BUSCO genes has been publicly released as a new tool, ChromSyn, under a GNU General Public License v.3.0 and is available on GitHub (https://github.com/slimsuite/chromsyn). Synteny of the hapA and hapB v3 *A. psidii* genomes was analyzed with ChromSyn v0.9.3 and further comparisons were also made against the fully phased, chromosome-level wheat leaf rust pathogen genome, *Puccinia graminis* f. sp. *tritici* (*Pgt 21-0*) (75).

For the nucleotide synteny analysis chromosomes of hapA and hapB v3 were aligned with Minimap2 v.2.28 (with options: --cs -cx asm20 --secondary=yes) (60). Matches with at least 80% identity and 1 kbp in length were used for generating a dotplot with Matplotlib v.3.9.1 (76).

To perform gene based synteny analysis, orthologs between haplotypes were identified with blastp v.2.15.0 (72, 73). Matches were filtered to ensure a minimum of 50 % sequence coverage for both query and reference sequences, along with at least 70 % identity. TE density and 5mCpG/CpG ratio were calculated using non-overlapping 10 kbp windows, with GenomicRanges v.1.54.1 (77). The results were visualized with karyoploteR v.3.17 (78).

### 2.8 Dinucleotide and DNA methylation analysis

For calculating observation/expectation (O/E) ratio of dinucleotides, the following formula was used to calculate the ratio for each chromosome separately (79):

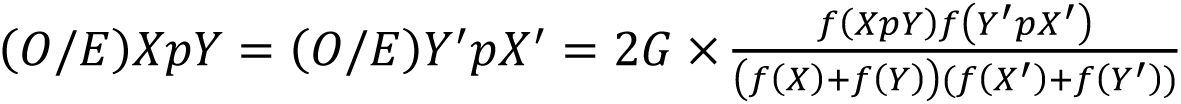

Where XpY represents the dinucleotide in one strand, Y’pX’ represents the complementary dinucleotide in the opposite strand, and G represents the length of the chromosome. HiFi reads with kinetics were generated from raw subreads by pbccs (v.6.4.0, https://github.com/PacificBiosciences/ccs) with the ‘—hifi-kinetics’ tag. Jasmine v.2.0.0 was applied to predict 5mC of each CpG site from PacBio HiFi reads, and pbmm (v.2 1.14.99, https://github.com/PacificBiosciences/pbmm2.git) was then used for aligning HiFi reads to the genome. Pb-CpG-tools (v.2.3.2, https://github.com/PacificBiosciences/pb-CpG-tools.git) was used for generating site methylation probabilities from mapped HiFi reads; only sites that have at least a 50-modification score were regarded as methylated fractions.

The total number of methylated sites within non-overlapping 1 kb windows across the genome, was calculated as:

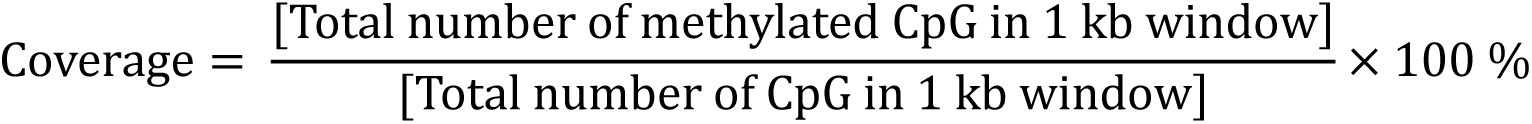

For confirming the 5mC methylated fractions, 5mC base calling and mapping were also performed using Dorado (v0.7.3+, https://github.com/nanoporetech/dorado) with the sup DNA model v.4.1.0.

Statistical differences between haplotype A and haplotype B were analysed using Student’s t-test by applying the SciPy module ttest_ind (80). Correlations between TE identity and GC content, as well as between TE identity and counts of CpG fractions per kbp, were analyzed using Pearson’s correlation coefficient, applying the SciPy modules pearson (80).

For examining whether transposons were hypermethylated, a permutation test was performed to compare random occurrences of 5mCpG fractions and observed 5mCpG fractions of TEs, coding sequences (CDS), and untranslated regions (UTRs). Random shuffling of 5mCpG fractions across CpG dinucleotides was performed using bedtools shuffle v.2.31.0 (81). The shuffling was constrained to occur only at CpG sites using the following parameters: -chrom (restrict shuffling to the same chromosome), - noOverlapping (prevent overlapping features), and -incl (specify CpG site locations). Random seeds were generated using the random module of Python v.3.12, and the shuffling process was repeated for 1,000 iterations. Percentage of 5mCpG fractions occurred in TEs, CDS and UTRs were calculated with pybedtools v.0.10.0 (82), the two-sided p-value was calculated by comparing simulation and observation, visualized with seaborn v.0.13.2 (83).

### 2.9 Centromere analysis

Contact maps visualized with Juicebox software (v.2.17.00) (62, 63) were used to estimate the position of centromeres. Statistical differences of TE coverage and 5mCpG/CpG ratio between centromeric and non-centromeric regions were performed using paired sample t-Test by applying the SciPy module ttest_rel (80).

### 2.10 Mating type analysis

To identify *MAT* genes, the *bW-HD1*/*bE-HD2* and *PRA* genes of *Pt BBBD* isolate (42) were used as queries in a blastp v.2.15.0 (72, 73) search. Following the confirmation of the PRA genes position, all Open Reading Frames (ORFs) ranging from 90 bp to 300 bp that contained ‘CAAX’ motifs were considered potential pheromone peptide candidates. Tandem repeat sequences were manually assessed for each candidate. Sequences less than 210 bp encoding at least two tandem repeats were classified as mfa1. Sequences less than 120 bp with ’CAAX-Stop’ motifs but lacking tandem repeats were classified as MFA2. Sequences greater than 210 base pairs containing tandem repeats like those in MFA1 were categorized as MFA3, based on observations in *P. graminis* f. sp. *tritici*.

A specific pattern ‘[Q/E]WGNGSH[X]C’ (where X can be any amino acid), was frequently found in the candidate mfa1/3 sequences, consistent with reported mfa1/3 in cereal rust fungi (38, 42). This pattern was used for further filtering candidates. To predict mfa2, the ‘CAAX-Stop’ motif was employed to capture ORFs ranging from 30 to 40 amino acids. CDS sequences of the identified mfa candidates were searched across the with blastn v.2.15.0 (72, 73) to identify orthologs, with all hits filtered as described above. The amino acids of the candidate mfas were aligned with ClustalW v.2.1 (84), and sequence logo graphs were generated using PlotnineSeqSuit (85).

To investigate the structure of the mating-type locus, the same approach used by Luo et al. (2024) for detecting genetic degeneration was employed. Proteinortho v.6.0.22 (86) was used to identify one-to-one orthologous gene pairs with on sister chromosomes, applying the -synteny and -singles tags. For each gene pair, coding-sequence-based protein alignments were generated with Muscle v.3.8.1551 (87) with default settings. Subsequently, PAML v.4.9 (88) was used to calculate synonymous divergence values from each alignment. TE density, 5mCpG ratio, and gene density were calculated with GenomicRanges v.1.54.1 (76) in 10 kb sliding windows without overlapping. Results were visualized with karyoploteR v.3.17 (78).

### 2.11 Allele specific expression analysis

Raw reads obtained in 2.4 were inspected using FastQC v.0.12.1 (https://www.bioinformatics.babraham.ac.uk/projects/fastqc/). Fastp v.0.23.4 was used for adapter trimming (Chen et al., 2018). Trimmed reads were mapped to the merged hapA and hapB *A. psidii* v3 transcriptome using HISAT2 v.2.21 (89) with parameter ‘--no-unal’ so that only reads that mapped to the *A. psidii* transcriptome were retained for further analysis. Salmon v.0.13.1 (90) was used to quantify *A. psidii* transcripts from the reads aligned to the transcriptome. Transcript level counts were imported into R v.4.4.1 (91) and summarized to gene level counts using the tximport package v.1.32.0 (92).

Allele specific differential gene expression analysis was carried out to identify which *A. psidii* genes were significantly upregulated *in planta* compared to urediniospores. The edgeR v.4.2.2 package in R (93) was used as it allows comparisons between multiple treatments in a single analysis. Genes that were lowly expressed were filtered out using ‘filterbyExpr’ function in R. The remaining gene counts were normalized with Trimmed Mean of Means (TMM) method using ‘calcNormFactors’ function in edgeR. The quality assessment and expression profiles of the samples were visualised using principal component analysis (PCA) and Pearson Correlation. This was achieved through the ‘plotMDS’ function from the Limma package v.3.60.6 (94), ‘cor’ function and pheatmap package v.1.0.12 (74) in R. To identify significantly upregulated genes in planta relative to urediniospores, pairwise comparisons between in planta time points and ungerminated urediniospores were made (2 dpi vs urediniospores, 3 dpi vs urediniospores, 4 dpi vs urediniospores, 5 dpi vs urediniospores, 6 dpi vs urediniospores). Genes with a Benjamini-Hochberg false discovery rate (FDR) of p < 0.05 and log-fold change > 2 or < -2 were considered biologically significant.

## Data availability

Genome assembly and raw sequencing data have been deposited in NCBI under BioProject accession PRJNA810572 and PRJNA810573. The functional annotations and TE annotations are publicly available through Zenodo (DOI: 10.5281/zenodo.15522519).

## 3. Results

### 3.1 *Austropuccinia psidii* displays uniform infection dynamics on host species separated by 65 million years of evolution

To understand the infection dynamics of the pandemic biotype of *A. psidii* on susceptible host species across the Myrtaceae family we selected four species, *S. jambos*, *M. quinquenervia*, *M. alternifolia* and *L. scoparium*, that are widely distributed across the phylogeny and separated by ∼65 million years of evolution (Fig. 1A) (95).

**Figure 1:**
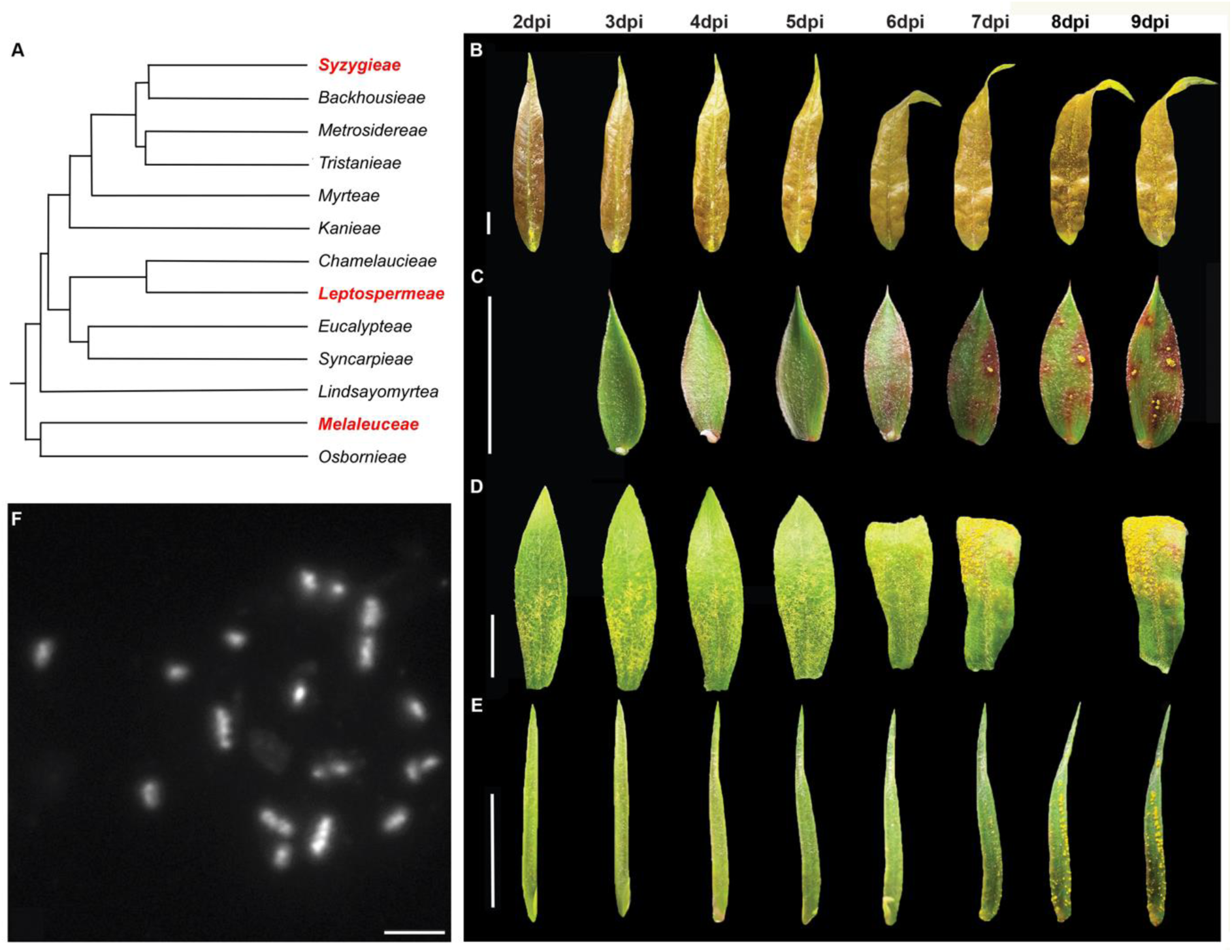
The pandemic biotype of *A. psidii* displays uniform infection dynamics across distantly related plant species. (A) Topology of genera within the Myrtaceae family. Genera that have species used in the infection assay are highlighted in red. The figure was adapted from Biffin et al. (2010) and Grattapaglia et al. (2012). (B-E) Infection symptoms developed between 2-/3-and 9-days post inoculation (dpi) for (B) *Syzygium jambos*, (C) *Leptospermum scoparium*, (D) *Melaleuca quinquenervia* and (E) *Melaleuca alternifolia*. Scale bars represent 1 cm. (F) Cytological analysis of *Austropuccinia psidii* chromosomes. Chromosomes are counterstained with 4’,6-diamidino-2-phenylindole (DAPI). They form a bimodal karyotype, with large chromosomes featuring clear primary constriction and two chromosome arms, and small chromosomes lacking visible primary constrictions. Scale bar: 2 μm.

Whole-plant infection assays revealed very similar symptom development across all four species when incubated under the same conditions (Fig. 1B-E). We did not observe any macroscopic myrtle rust disease symptoms until six days post infection (dpi), when the first lesions appeared, characterized by yellow or reddish spots and leaf curling. By seven dpi, the initial sporulating pustules emerged, with their numbers increasing steadily until the end of the experiment at nine dpi (Fig. 1B-E, Fig. S1, S2).

### 3.2 Cytology and near-T2T genome analyses reveal karyotype conservation and inter nuclear chromosome transfer in *A. psidii*

To characterize chromosome size distribution and number in *A. psidii*, we conducted a cytological study. This revealed that *A. psidii* has chromosomes belonging to two distinct size classes. In a haploid mitotic metaphase cell, chromosomes were organized into a bimodal karyotype comprising twelve larger and six smaller chromosomes (Fig. 1F). The twelve large chromosomes displayed clear primary constrictions and two chromosome arms, while the six smaller chromosomes lacked visible primary constrictions and were considerably shorter (Fig. S3).

We followed up on these cytological karyotype estimations by generating the first near-telomere-to-telomere (near-T2T) genome for *A. psidii*, because the previously published *A. psidii* genome assembly was fragmented, with over 66 scaffolds per haplotype and only three complete chromosomes (Table 1) (21). This limited the genome biology insights obtained from this partial phased assembly, which contains significant residual assembly errors. To obtain a fully phased near-T2T assembly of the *A. psidii* pandemic biotype, we combined HiFi and ONT long reads, with Hi-C data, followed by additional Hi-C scaffolding (Table S1). This approach yielded two haploid assemblies of approximately 1 Gbp each, which we named hapA and hapB (Table 1).

**Table 1:** T2T genome assembly statistics of *Austropuccinia psidii* v3. Basic genome statistics comparing the v1 *Austropuccinia psidii* genome (Tobias et al., 2021), against the current genome assemblies (v3).

| Genome | # Scaffold (contig) | # Chr | Total Length (bp) | Max Length (bp) | N50 (bp) | L50 | GC (%) | BUSCO (%)* |
| --- | --- | --- | --- | --- | --- | --- | --- | --- |
| hap1_v1 | 66 (3,817) | 3 | 1,018,398,822 | 124,273,364 | 56,243,252 | 7 | 33.8 | C: 88.1 [S: 78.6; D: 9.5] |
| hapA_v3* | - | 19 | 1,073,490,195 | 85,591,751 | 60,211,033 | 8 | 33.84 | C: 91.2 [S: 90.3; D :0.9] |
| hap2_v1 | 67 (8,637) | - | 934,744,333 | 89,073,602 | 52,409,407 | 7 | 34.26 | C: 70.3 [S: 67.0; D: 3.3] |
| hapB_v3* | - | 17 | 982,558,553 | 84,687,478 | 60,506,416 | 7 | 33.86 | C: 90.9 [S: 90.0; D: 0.9] |
| Dikarion_v3 | (270) | 36 | 2,079,842,500 | 85,591,751 | 60,211,033 | 15 | 33..86 | C: 92.0 [S: 2.8; D: 89.2] |

\*Complete BUSCOs (C); Complete and single-copy BUSCOs (S); Complete and Duplicated BUSCOs (D).
\*To compute BUSCO value, one chr14 is distributed into hapB.

The assembled genome indicates a haploid karyotype of 18 chromosomes of varying sizes, ranging from ∼86 Mb to ∼28 Mb per chromosome. Surprisingly, Hi-C contact maps (Fig. 2) revealed that hapA contained 19 and hapB 17 chromosome-sized scaffolds, suggesting that chromosome chr14B relocated to the hapA nucleus and is now stably inherited via urediniospores through generations. We ordered and numbered scaffolds by size, hence scaffold one is the largest of ∼86 Mb and scaffold 18 is the smallest of ∼28 Mb. Fourteen hapA and 15 hapB scaffolds are telomere to telomere (T2T), indicating likely full-length chromosomes (Table 1 and Fig. 2). The other five scaffolds in hapA and two scaffolds in hapB have a telomere at one end only. Two-thirds (24/36) of the annotated telomeres range between 9 – 12 kb (Table S2). The overall BUSCO gene completeness of 92 % is a slight improvement over 91.2 % for v1. The total genome size for v3 is comparable to v1 (Table 1).

**Figure 2:**
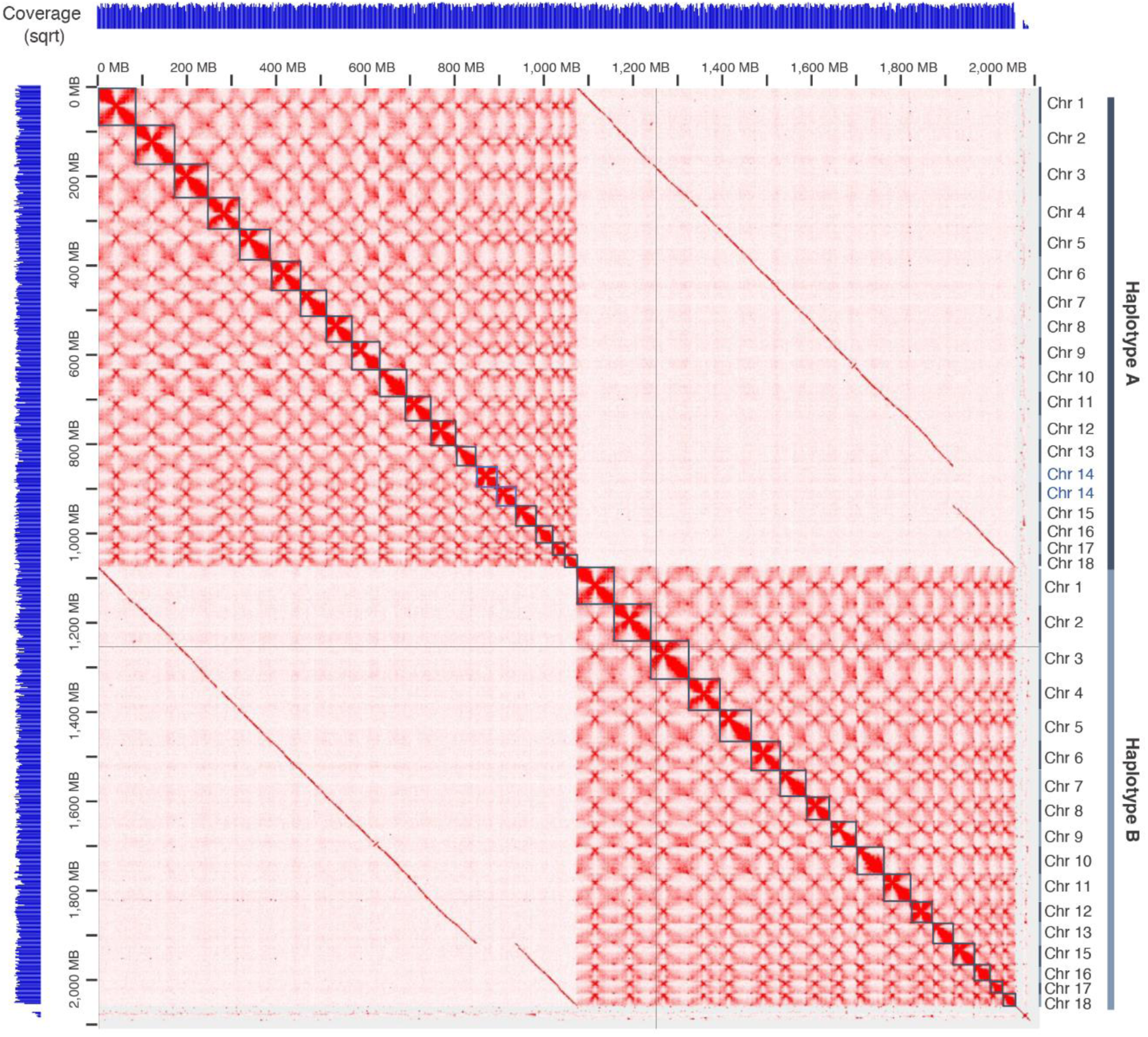
3D genome confirmation capture reveals inter nuclear chromosome transfer in *Austropuccinia psidii* between its two haploid nuclei. 3D genome interaction heat map of the 36 chromosomes of the *A. psidii* v3 genome visualised in Juicebox. Hi-C coverage indicated by pale blue bars, left and top. Black squares indicate the predicted 19 chromosomes in order of size in haplotype A and 17 chromosomes in haplotype B.

### 3.3 Gene synteny of conserved genes (BUSCOs) is mostly maintained when compared to distantly related rust fungus *Puccinia graminis* f. sp. *tritici* despite significant expanded genome size due to TEs

First, we confirmed the transposable element expansion reported for the *A. psidii* v1 genome assembly (21). Approximately 89.14 % of the near-T2T genome assembly of *A. psidii* is comprised of repetitive elements. Among these, Class I TEs: LTR, *Ty3* has the highest proportion across all chromosomes, followed by Class II: TIR: CACTA (Fig. S4). Then we tested the gene synteny between the two haplotypes of *A. psidii*. Overall, a conserved syntenic relationship was observed across most chromosomes (Fig. S5-S6), except for a large chromosomal inversion on chr7. Additionally, chr14 showed substantial structural variation, marked by multiple translocation events and gene duplications (Fig. S5-S6).

Since *A. psidii* and other known rust fungi share an identical karyotype, we investigated the conservation of the synteny of BUSCO genes. The synteny of BUSCO genes was compared between the two *A. psidii* haplotypes and to the chromosome assembly of *P. graminis* f. sp. *tritici* 21-0 (*Pgt* 21-0) (75). For this analysis, we developed ChromSync, a tool designed for the rapid identification of syntenic blocks based on BUSCO gene order. The synteny of BUSCO genes is largely conserved between the two *A. psidii* haplotypes with hardly any structural translocation of BUSCO genes amongst chromosomes of the two haplotypes. Similarly, the overall synteny of BUSCO genes was mostly conserved between *A. psidii* and *P. graminis* f. sp. *tritici* 21-0 chromosomes except for multiple rearrangements involving chr14 and chr2 of *A. psidii* and chr2 and chr11 of *P. graminis* f. sp. *tritici* 21-0 (Fig. 3).

**Figure 3:**
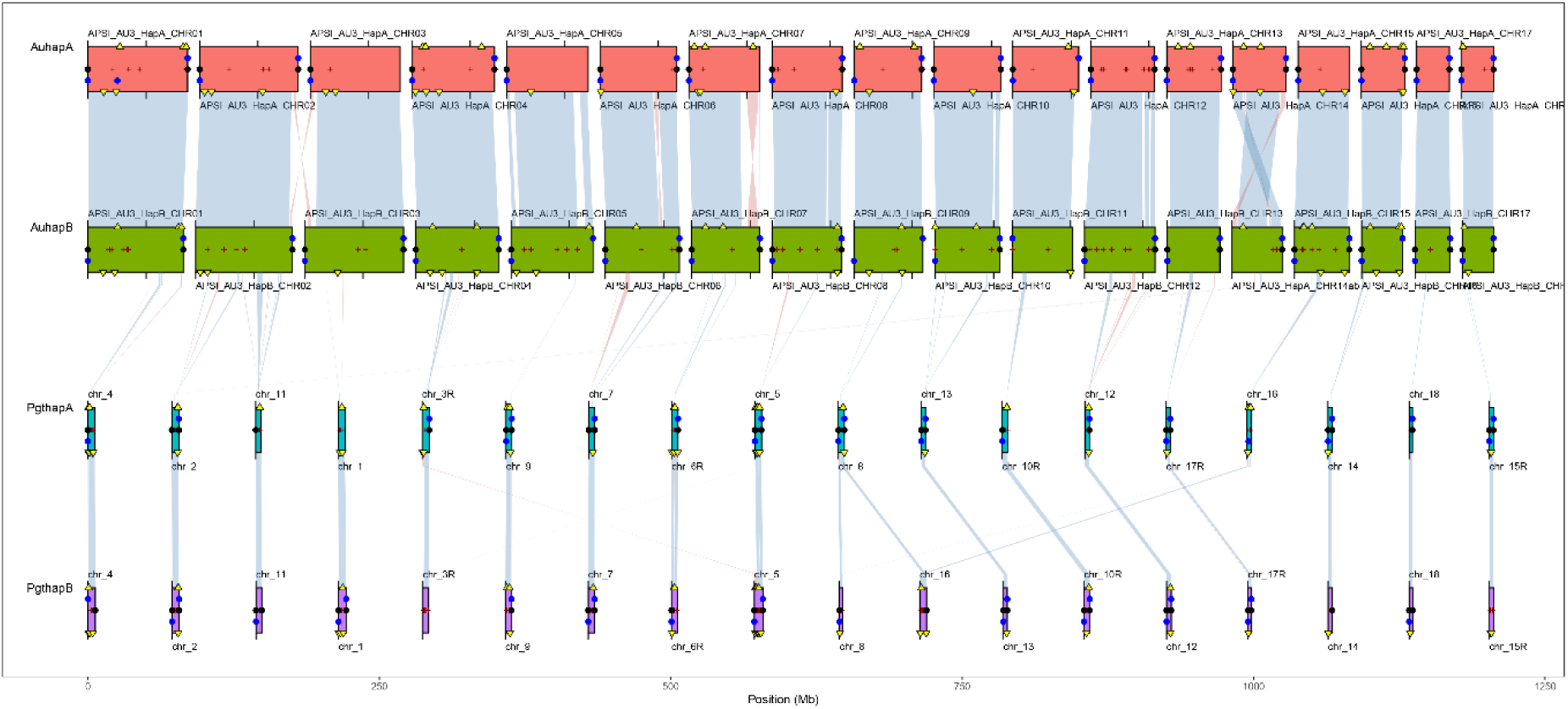
BUSCO gene synteny is larger conserved between *Austropuccinia psidii and Puccinia graminis* f. sp. *tritici*. The figures show ChromSyn BUSCO-derived synteny plots. Coloured blocks represent assembly scaffolds over 1 Mbp for each genome. Synteny blocks of collinear “Complete” BUSCO genes link scaffolds from adjacent assemblies: blue, same strand; red, inverse strand. Filled circles mark telomere predictions from Diploidocus (black) and TIDK (blue). Assembly gaps are marked as dark red +/- signs. Genes on the forward strand are marked above the centre-line, with reverse strand genes below. Pairs of phased haploid genomes for *Austropuccinia psidii* v3 (AuhapA and AuhapB) and *Puccinia graminis* f. sp. *tritici* 21-0 (PgthapA and PgthapB) are shown (75).

### 3.4 Detailed analysis of the *Austropuccinia psidii* near-T2T genome reveals specific DNA methylation patterns, CpG depletion via historic 5mC deamination, centromeres and a significantly extended mating-type PR locus

Using the new near-T2T genome assembly we extended the genome biology analyses of *A. psidii*, focusing on aspects that previously were intractable including CpG DNA methylation, centromeres, and detailed analyses of the mating-type loci. Consistent with previous studies, younger TEs (with >90 % family-level sequence identity) exhibit higher GC content and more available CpG methylation sites (SFig. 7). Both measures were also positively correlated with TE family age when measured as average percentage identity of individual TE insertions relative to the TE family’s consensus sequence (family-level sequence identity) (Pearson’s r = 0.18 and r = 0.12 respectively, both p < 1 × 10-16).

We also examined the genome wide dinucleotide composition. The genome-wide dinucleotide observed/expected (O/E) ratios show that CpG, ApC (GpT) and TpA are underrepresented, while CpC (GpG) is overrepresented (Fig. 4A). We determined the 5mCpG methylation levels on the remaining CpG sites and revealed that the ratio of methylated CpG sites varies across chromosomes from 0.37 to 0.54 (Fig. 4B). Chromosomes contained in hapB displayed a significantly higher ratio of methylated CpG sites compared to hapA (p < 0.005) (Fig. 4B). To investigate whether TEs were preferentially methylated, a permutation test of random occurrence of 5mCpG fractions was performed. The non-random occurrence of 5mCpGs suggest the TEs were hypermethylated (p < 0.01) whereas CDS (p < 0.01), UTRs (both p < 0.01) are hypomethylated (Fig. 4C). When focusing on TEs, the results revealed that younger TE families displayed significantly higher 5mCpG ratios than older TE families. This suggests that DNA methylation levels play a more important role in TE silencing of younger TE families.

**Figure 4:**
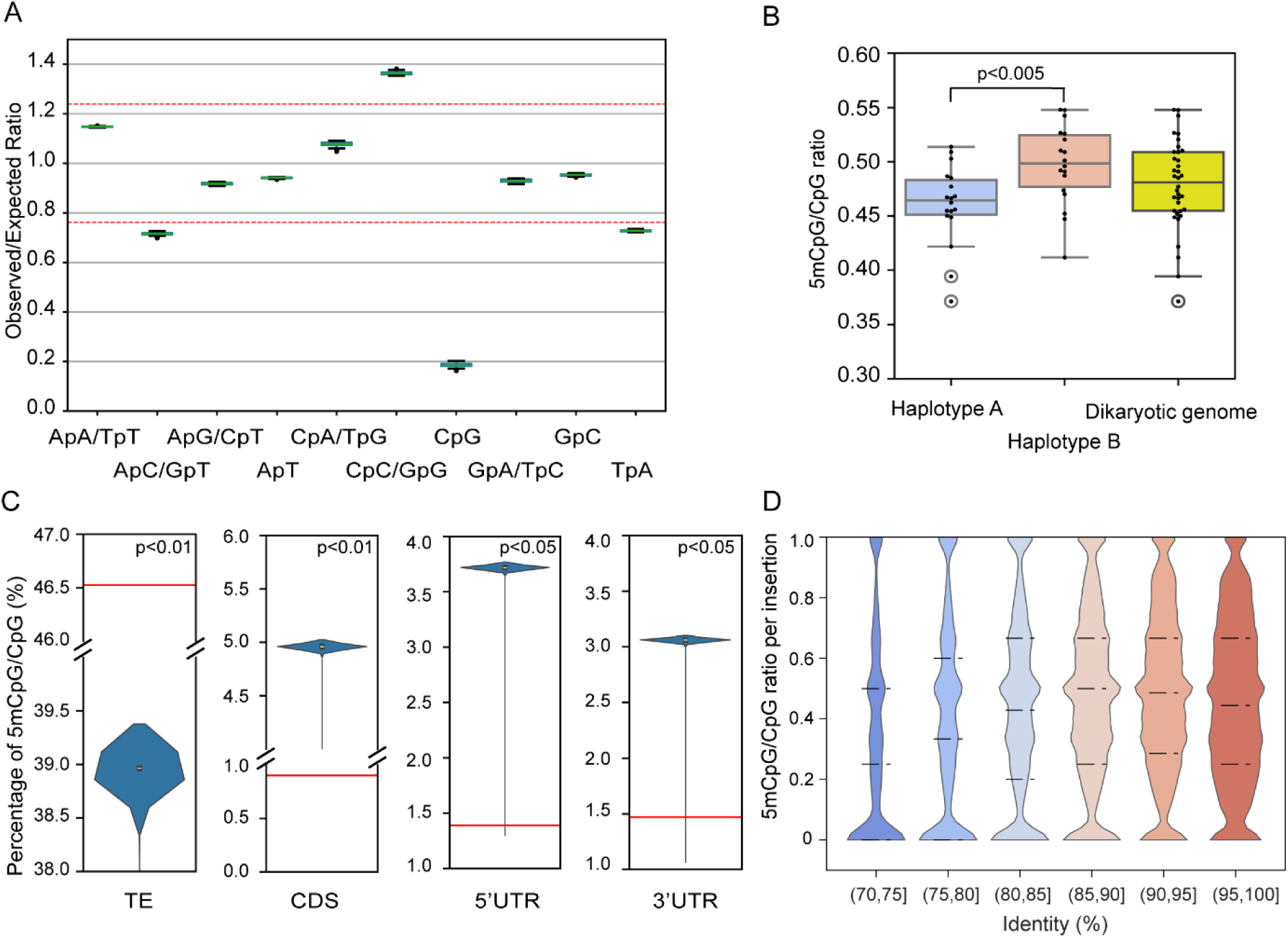
5mCpG methylation is unequally distributed across the *Austropuccinia psidii* genome and inversely corelated with TE age. (A) Dinucleotide observation/expectation (O/E) ratios across chromosomes. Red broken lines at 0.75 and 1.25 represent the threshold of underrepresentation and overrepresentation respectively. (B) Ratio of 5mCpG/CpG across chromosomes of haplotype A (blue), haplotype B (orange), and the diploid genome (yellow). The difference between haplotype A and haplotype B is statistically significant (p < 0.005). (C) Percentage of 5mCpG/CpG overlapping transposable elements (TEs), coding DNA sequence (CDS), 5’ untranslated region (UTR), and 3’ UTR in permutation test respectively, red lines represent observation in genome. (D) Ratio of 5mCpG/CpG per insertion of TEs. TEs are categorized into bins based on their sequence identity, with each bin representing a 5 % interval ranging from 70 % to 100 % identity. Dashed lines indicate the 25th, 50th and 75th percentiles, respectively.

Next, the centromeric regions of the repeat rich genome of *A. psidii* were investigated, guided on the classic bowtie structure clearly discernible in the Hi-C contact map obtained from the urediniospore stage (Fig. 2). Analysis of the nucleotide sequence conservation in centromeres between sister chromosomes showed that nearly all sister chromosomes maintained conserved synteny, except chromosome 8 (Fig. 5A). While the centromeric regions were conserved, their flanking areas contained syntenic gaps (Fig. 5A). Given that fungal centromeres are characterized by high TE coverage and cytosine hypermethylation – both important for stability – we analysed TE density and 5mCpG/CpG ratios. Centromeres showed significantly higher 5mCpG/CpG ratios (p < 0.01), but lower TE density compared to non-centromeric regions (p < 0.01) (Fig. 5B-C, SFig. 8).

**Figure 5:**
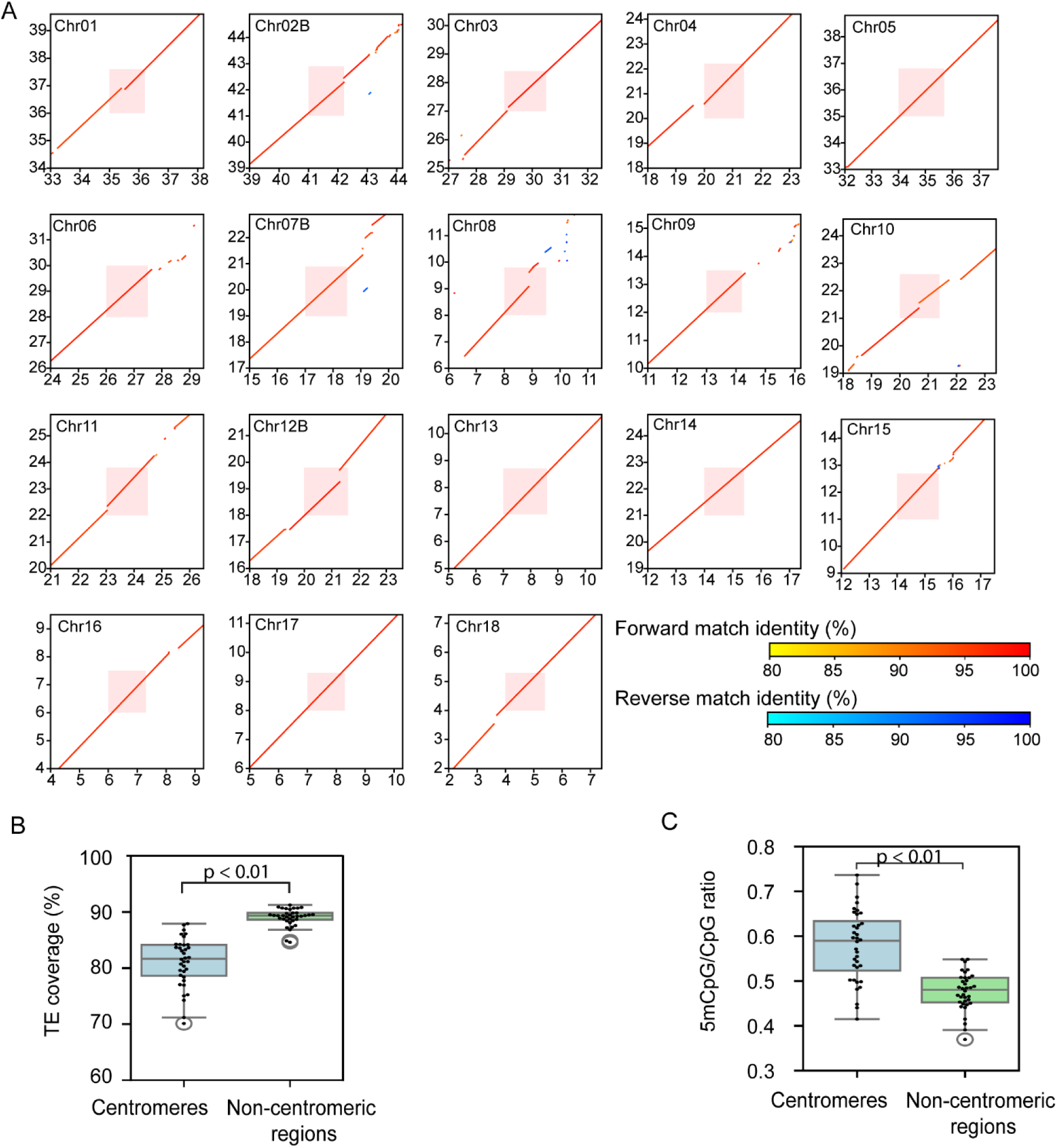
Centromeres are syntenic between orthologues chromosome pairs and hypermethylated. (A) Dot plot shows similarity of centromeres and their adjacent regions between haplotypes A and B. The x-axis represents chromosomes phased in haplotype B, while the y-axis represents chromosomes phased in haplotype A. Red dots represent regions aligned in the same orientation, while blue dots represent regions aligned in the reverse direction. The positions of the centromeres are highlighted in red. (B-C) Comparison of TE coverage (B) and the 5mCpG/CpG ratio (C) within centromeres and non-centromeric regions. Centromeres showed significantly higher 5mCpG/CpG ratios (p < 0.01) (B), whereas lower TE density compared to non-centromeric regions (p < 0.01) (C). Each dot represents the value on a single chromosome.

The genome biology of the two mating type loci (HD and PR locus) in the new near-T2T genome was also characterized. The HD locus encoding the two homeodomain transcription factors (*bW-HD1* and *bE-HD2*) is located on chr1. The PR locus encoding *Pra* pheromone receptor alleles (*STE3.2-2* and *STE3.2-3*) is located on chr5. The overall synteny of the HD locus was conserved between chr1A and chr1B, with no obvious changes in heterozygosity, gene order and TE density when compared to adjacent regions on the same chromosome (Fig. 6A and Fig. S9). This is in stark contrast to the PR locus, which is heavily enriched in TEs, depleted in genes, and extends over ∼10 Mbp (Fig. 6B-C). The 5mCpG DNA methylation levels do not appear to be elevated at the PR locus. The two *Pra* alleles appear to be either distal (*STE3.2-2*) or proximal to the centromeric region (*STE3.2-3*).

**Figure 6:**
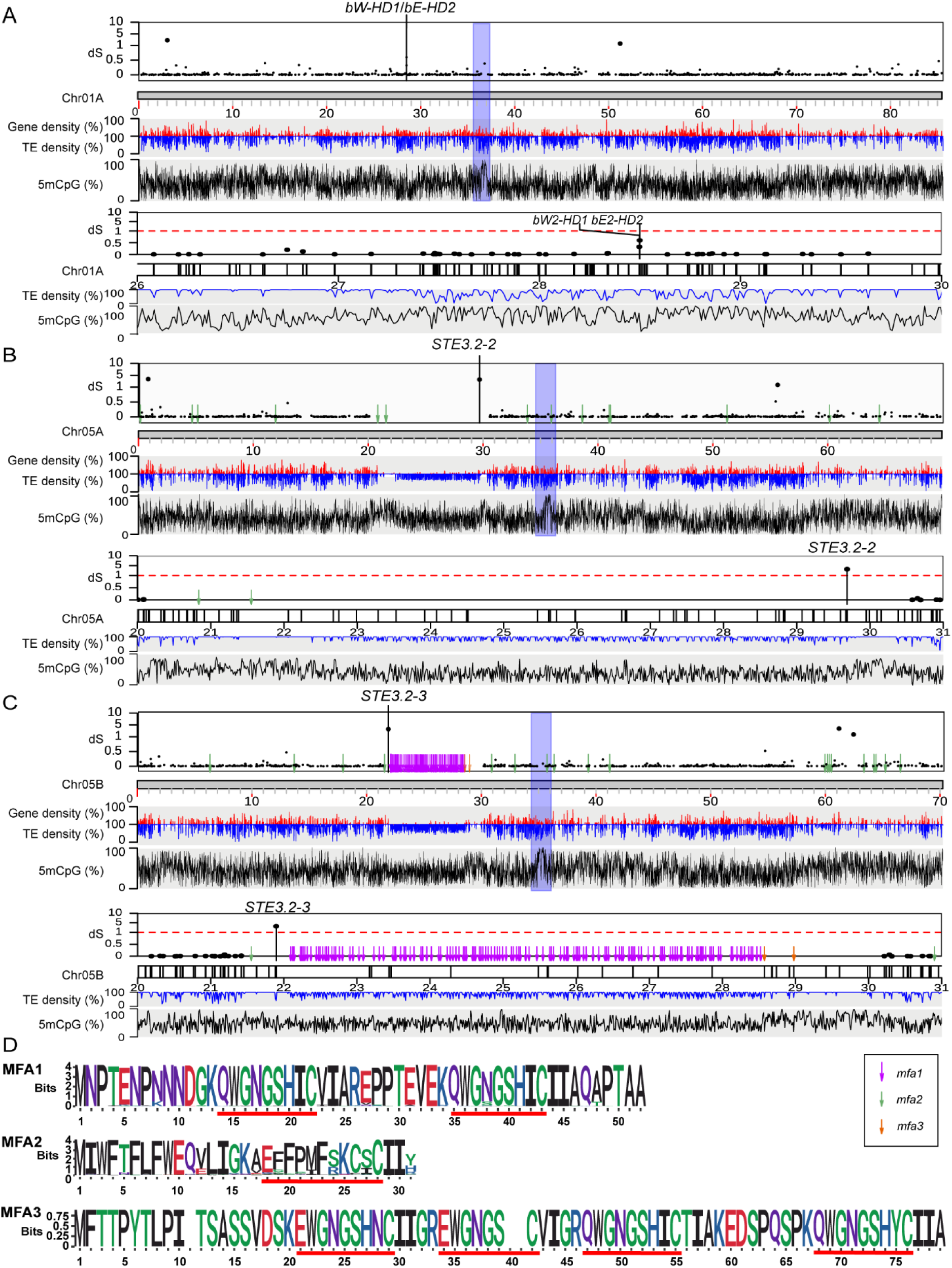
The PR locus is highly enriched in TEs and displays a proliferation of potential mating type pheromone peptide precursor genes. Synonymous divergence values (d_S_) for all allele pairs are plotted along chromosomes (A) 1A, (B) 5A and (C) 5B. In each panel, the top track shows the d_S_ values of allele pairs along chromosomes, where each dot on the top track represents d_S_ of a single allele pair. The second, third and fourth track show the average transposable elements (“TE”), gene (“gene”) density and ratio of 5mCpG/CpG (“5mCpG”) along chromosomes in 10 kbp-sized windows respectively. Predicted centromeric regions are marked with blue shading, predicted mfas are labelled with pink arrows (*mfa1*), blue arrows (*mfa2*) and orange arrows (*mfa3*). The lower tracks provide a detailed zoomed-in view of regions around (A) the HD locus and (B-C) PR locus, black markers on chromosome tracks represent genes. (D) Sequence logos of predicted MFA1, MFA2, and MFA3, with predicted mature peptide pheromone sequences indicated by red underlines. Amino acids are coloured based on chemical polarity.

Next, we identified all potential pheromone peptides recognized by the two *Pra* receptors. To identify *mfas* in *A. psidii*, we screened ORFs between 90 and 300 base pairs encoding peptides contained ’CAAX’ motifs. The MFA candidates were classified into three categories based on specific sequence features and numbers of tandem repeats: *mfa1*, *mfa2* and *mfa3*. Manual inspection of tandem repeats revealed the sequence pattern ‘[Q/E]WGNGSH[X]C’ with high frequency among most potential *mfa*1/3s, therefore, this pattern was used for further filtering *mfa*1/3s (Fig. 6D).

In total, 14 copies of *mfa2* on chr5A, 19 copies of *mfa2*, 137 copies of *mfa1* and 4 copies of *mfa3* on chr5B were identified (Fig. 6D). All *mfa1* and *mfa3* clustered between 22 Mb-29 Mb of chr5B, which is likely to be the recombination suppressed region of the PR locus containing *STE3.2-3* (Fig. 6C). In contrast, *mfa2* genes were dispersed across both chr5A and chr5B, showing no obvious clustering pattern. Next, MFA coding sequences were further searched in the whole genome and identified an additional 314 copies of *mfa2*s on non-PR locus containing chromosomes.

### 3.5 Transcriptional regulation reveals two waves of expression during early *vs* late plant infection and includes allele specific expression of candidate effectors

Illumina RNA sequencing was used to understand gene expression of *A. psidii* during infection of *S. jambos* over time. The experimental design included ungerminated urediniospores and 2-6 days post infection on *S. jambos* (Table S3).

We confirmed the accuracy of the RNA-seq analysis and relatedness of biological replicates, using PCA cluster analysis. The analysis suggests that the differential gene expression patterns were strongly linked to sampling time, showing that replicates of earlier timepoints (2, 3, and 4 dpi) clustered separately from later timepoints (5 and 6 dpi), and ungerminated urediniospores (0 dpi) (Fig. 7B). This suggests low biological variability among replicates and high degree of similarity in their expression profiles. The first principal component accounts for 84 % of the variation between time points, and the second component accounts for 9 %.

**Figure 7:**
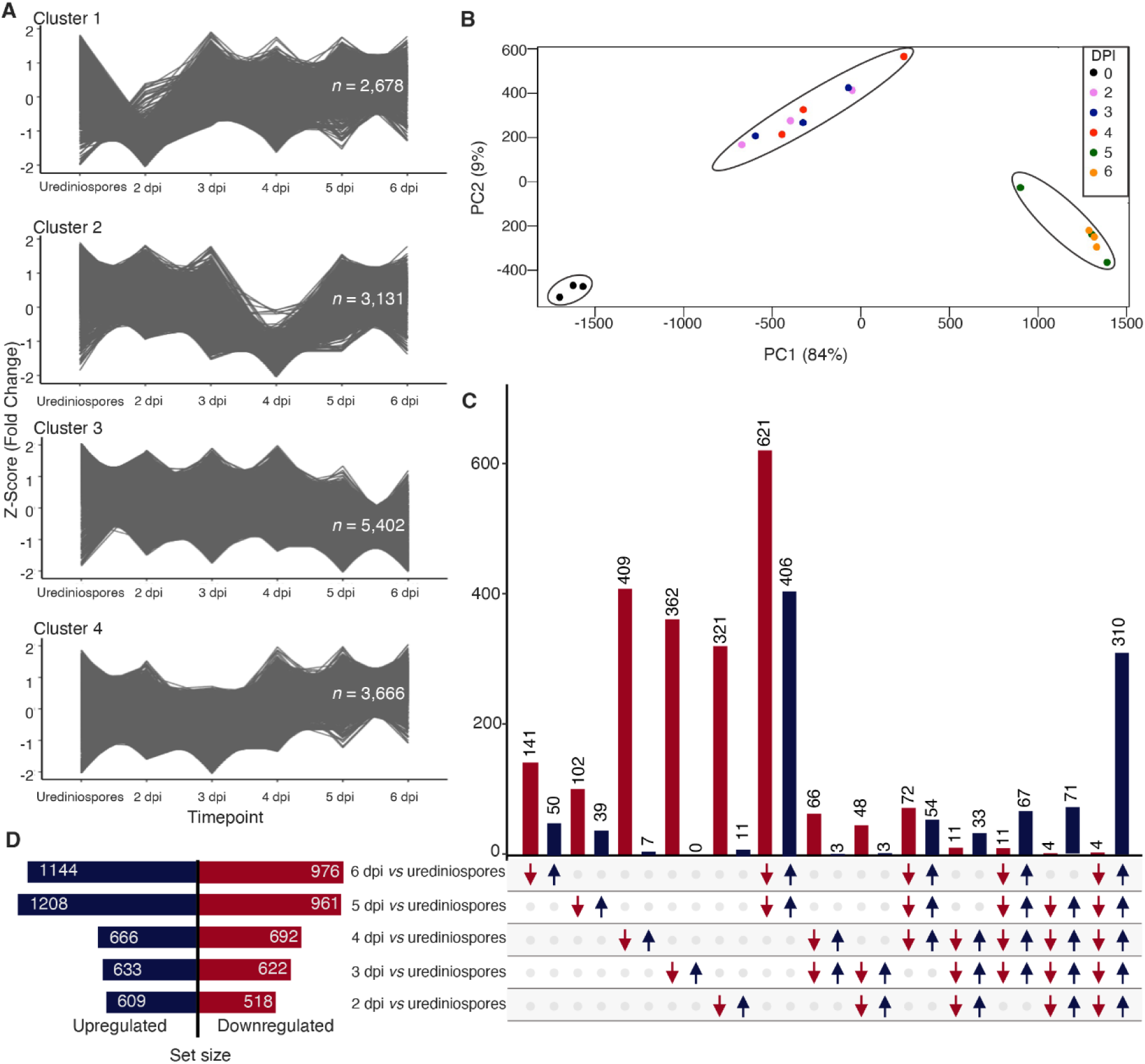
*Austropuccinia psidii* displays a biphasic transcriptomic reprograming upon infection of *Syzygium jambos* over time. (A) *k-*means cluster plots from RNA-seq data analyzed using the TCseq package, utilizing the fold change (FC) z-score option for normalization. Each cluster represents distinct expression patterns across the time points. The number of genes that belong to each cluster are highlighted inside the graph; (B) Multidimensional Scaling (MDS) plot illustrating the relationships among transcript expressing profiles from six different sample groups; (C) UpsetR plot showing differentially expressed genes specific to each pairwise comparison. The number of up regulated (dark blue bars, arrow directed upward) and down regulated (red bars, arrow directed downward) transcripts are presented on the upper side of each intersection bar. (D) total changes in the *A. psidii* expression profile *in planta* when compared to resting urediniospores. The total number of differentially expressed genes (DEGs) for haplotype A and B, in each pairwise comparison is represented by dark blue (up regulated) and red (down regulated) bars.

To identify important pathogenicity components in the expression profile of *A. psidii* during infection, allele specific expression analysis was performed between each timepoint of infection (2, 3, 4, 5, and 6) in *S. jambos* and the ungerminated urediniospores (0 dpi). In total, 3,767 differentially expressed genes (DEGs) were identified (Table S4), of which 1,366 were upregulated in at least one time point *in planta* relative to ungerminated urediniospores (Fig. 7C-D). More specific overlapping DEGs were identified in latter time-points (5 and 6 dpi) when compared to earlier time-points (2, 3 and 4 dpi) (Fig. 7C-D). Since the focus was the molecular profile of *A. psidii* during interaction with its host, the remainder of this study focused on DEGs that were upregulated *in planta*. A total of 310 genes were commonly upregulated across all timepoints, suggesting a common repertoire throughout infection, while 442 genes were upregulated only at 5 and 6 dpi, and 34 genes were commonly upregulated at 2, 3 and 4 dpi (Fig. 7C, Table S4). The top upregulated genes in all timepoints were identified from the DEGs table and resulted in 22 genes, 16 of those, classified as candidate effectors (CEs) (Table 2).

**Table 2.**
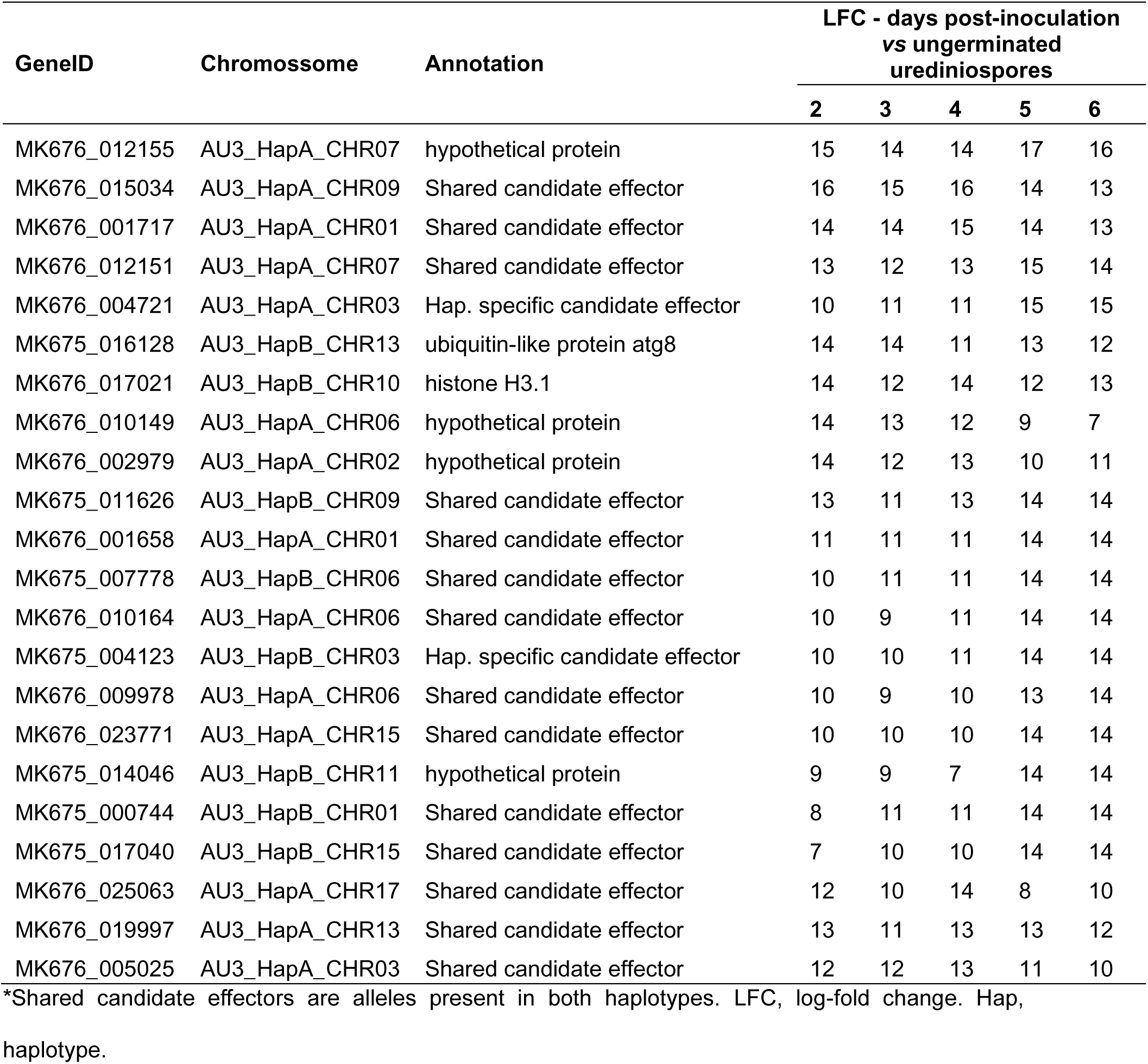
Candidate effectors are enriched in highly expressed genes *in planta*. Top upregulated genes *in planta* compared to ungerminated urediniospores.

**Table 3.** Predicted localisation of candidate effectors Allelic and hemizygous candidate effectors (CEs) of *Austropuccinia psidii* v3 genome, haplotype A (hapA) and haplotype B (hapB). Ext – extracellular, PM – Plasma Membrane, Cyt – cytosolic

| Haplotype | # Secreted CEs | Shared |  |  |  | Hemizygous |  |  |  |
| --- | --- | --- | --- | --- | --- | --- | --- | --- | --- |
|  |  | # CEs | # Ext. | # PM | # Cyt. | # CEs | # Ext. | # PM | # Cyt. |
| hapA | 1206 / 283 | 253 | 190 | 13 | 50 | 30 | 23 | 2 | 5 |
| hapB | 979 / 240 | 215 | 167 | 6 | 42 | 25 | 14 | 1 | 10 |

To obtain biological meaningful information from differential expression results, putative functions were assigned to genes significantly upregulated *in planta*. Forty-eight percent (657/1366) of all upregulated genes were annotated with GO terms, and/or PFAM terms (Table S4).

Plant pathogens typically use secreted effector proteins to facilitate host colonization (Baid et al., 2023). Considering the presence of two copies of Chr14 in hapA, for Au3, 1206 and 979 secreted proteins were identified for hapA and hapB.

From those, 283 (23.4 %) and 240 (24.51 %) are predicted to be candidate effectors based on the evidence of changes in expression during *S. jambos* infection. Most of the CEs were predicted to be secreted to extracellular spaces. Thirty and 25 CEs were hemizygous and only present in hapA or hapB including two in the top overexpressed genes *in planta* (Table 2-3). Notably, 87 % of the CEs have alleles present in both haplotypes. Among these shared alleles, only 32 display consistent expression patterns, while the remaining alleles exhibit allele-specific expression (Fig. 8, Table S5).

**Figure 8:**
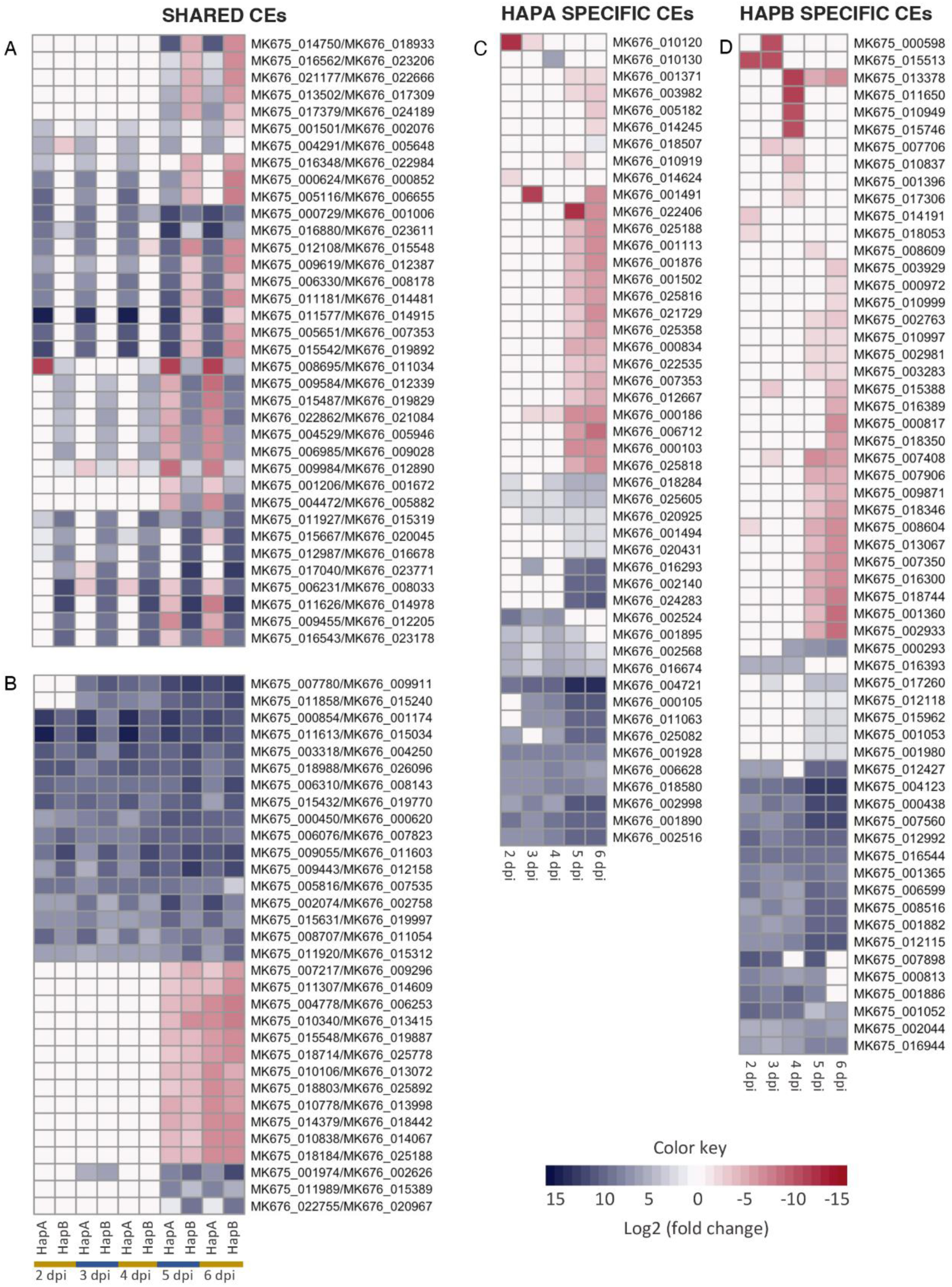
Haplotype specific expression of candidate effectors appears to be common in *Austropuccinia psidii*. Heatmaps of differentially expressed candidate effectors (CEs) *in planta* when compared to ungerminated urediniospores over time. Contrasting expression of alleles in haplotype A (hapA) and haplotype B (hapB) (A); alleles with the same pattern of expression in hapA and hapB (B). HapA specific CEs (C); hapB specific CEs (D). Days post-inoculation (dpi).

## 4. Discussion

The pandemic biotype of *A. psidii* exhibits an unusual lifestyle compared to other rust fungi, characterized by an expanding list of host species within the Myrtaceae (7). This pathogen’s dikaryotic nature, combined with its 1 Gb haploid genome size and high repeat content, presents significant challenges for genome assembly (21). These challenges can now be addressed through the integration of advanced sequencing technologies such as PacBio HiFi, ONT ultra-long reads and Hi-C data (23) permitting deeper investigation into the genome biology and adaptive evolutionary potential of this damaging pathogen. Here, biological approaches and the assembly of a fully phased genome for the pandemic lineage of *A. psidii* were integrated to answer important biological questions.

### 4.1 Karyotype conservation and genome plasticity in *A. psidii*

Our research shows that *A. psidii* has 36 dikaryotic (2n) chromosomes consistent with karyotypes of other distantly related rust fungal species (96, 97), and despite its phylogenetic placement within the family Sphaerophragmiaceae (8, 98). Of the assembled chromosomes, 29 have telomeres at both ends indicating complete chromosomes. Early karyotyping efforts underestimated chromosome numbers in rust fungi, reporting n = 3-6 (99, 100) due to small size and challenges in arresting the metaphase state in biotrophs. Subsequent studies on rust species, including *Melampsora lini* (96) and *P. graminis* f. sp. *tritici* (97), consistently identified a haploid chromosome number of *n* = 18. These findings have been recently further confirmed with new technology that has enabled phasing and scaffolding of genomes for *P. triticina* (24, 67), *P. striiformis* f. sp. *tritici* (26, 29), *P. polysora* f. sp*. zeae* (28), and *M. lini* (30). This suggests that the chromosome numbers are highly conserved among the rust fungi which diverged about 85 mya (98).

In *A. psidii*, the large genome and chromosome size are primarily attributed to the amplification of repetitive DNA sequences, including transposable elements (21). The large genome of *A. psidii* prompted our cytological investigations based on the expectation that chromosomes are likely to be more visible at metaphase state than in cereal rust fungi. Our cytological study found that *A. psidii* chromosomes can be clearly separated into two distinct size classes of 12 large and 6 small chromosomes, resulting in a bimodal karyotype, which is an interesting feature often seen in birds (98). This marks the first cytological study of *A. psidii* chromosomes, with the sizes aligning well to the distribution determined from sequencing analysis, suggesting that the karyotype reflects the fungal genomic structure. Further comparative genomic work among rust fungi covering the whole phylogeny will help to answer questions on karyotype conservation and genome evolution in response to continuous host adaptation.

### 4.2 Unequal chromosome numbers within the *A. psidii* haploid nuclei

Dikaryotic rust fungal evolution is characterized by extensive mitotic division leading to high levels of heterozygosity (24). The observation of uneven chromosome counts in two haplotypes have been previously reported in *P. graminis* f. sp. *tritici* 21-0, where both copies of chr11 were assigned to the same haplotype based on Hi-C analysis (75). Since the nuclei fusion is absent during asexual reproduction of rust fungi (101), this ‘chromosome-swap’ is likely to occur during meiosis. Mis-segregation events during this process can result in both copies of chr14 being directed to the same nucleus. Since mitosis lacks nuclei fusion, it is plausible that this chromosome swap can be inherited asexually. Our work shows greater homology between predicted proteins for chr14 compared to all the other chromosomes (Fig. S5). This greater homology may be explained by homologous recombination (102, 103) during extended non-sexual reproduction, whereby mutations are repaired based on the paired chromosome, while the other chromosomes, in separate nuclei, gain mutations and diverge. A speculative explanation is that a meiotic event in a precursor biotype to the pandemic developed this nuclei chromosome arrangement and asexual proliferation led to the dominant pandemic spore type that has expanded globally infecting a broader host range. The finding of a Brazilian biotype closely related to the pandemic *A. psidii* (13), may present a useful opportunity to further investigate this phenomenon.

An alternative explanation could involve a mis-segregation event during mitotic division followed by selection or random drift on isolates that carry unequal chromosome numbers in their nuclei.

Overall, it is noteworthy that chr14 of *A. psidii* and chr11 of *P. graminis* f. sp. *tritici* are not orthologous and that a recent study on two ascomycete fungal species also described unequal haploid chromosome numbers in bi-and multinucleated spores. Hence, it might be much more common to observe unequal haploid chromosome numbers in fungi that carry multiple nuclei in the same cytoplasm. Yet the evolutionary and adaptive implications are currently not known (104). It is unclear if this genome plasticity can fulfill a similar function to aneuploidy in yeast species, such as *Saccharomyces cerevisiae* and *Candida albicans*, where it can occur on any chromosome and contributes to adaptative phenotypes (105–107).

### 4.3 DNA methylation plays a role in transposable element silencing in *Austropuccinia psidii*

It is widely accepted that the CpG depletion of many animal genomes is caused by methylation and mutational loss of cytosines at CpG site (108–110). In many genomes, including the 1 Gb *A. psidii* genome compared to other rusts, there is a negative correlation between genome size and the O/E ratio of the CpG dinucleotides, suggesting that the suppression of TE activity by the 5mCpG deamination is essential for the their long-term accommodation of TE’s in the host genome (37).

We observed significant genome wide CpG depletion, likely to be the result of deaminated 5mCpG within TEs. Previous analysis on the v1 genome indicated that low GC content is unlikely to result from RIP mutation as *A. psidii* displayed insignificant RIP indices and an absence of genes associated with RIP mutation mechanisms. Hence the low GC content was hypothesised to be the consequence of genome wide deamination of methylated cytosines which cause C-to-T transition mutations (21). The current findings provide further support for this hypothesis, as 5mCpG/CpG ratio and GC content is higher in younger TE insertions than in aged insertions. This suggests that DNA methylation is the initial mechanism that silences younger TEs, and deamination and C-to-T are secondary TE silencing mechanisms (111). Additionally, the preferential cytosine methylation of TEs compared to genic regions of the genome highlights the role of DNA methylation in silencing these elements, contributing to overall genome stability.

Consistently, centromeres in rust fungi have been reported to be hypermethylated (29, 112), which is hypothesized to maintain chromatin stability during nuclear division and chromosome segregation (113). The identification of hypermethylated centromeres in *A. psidii* is consistent with these previous observations. Unlike other rust fungi, we did not observe TE coverage enrichment in centromeric regions compared to non-centromeric regions. It is currently unclear if this is a biological feature or a technical artifact. It could be that we missed centromeric TEs by employing the previously generated *de novo* TE database based on genome v1, which might have lacked centromeric sequences (21). Yet careful follow up investigation of centromeric regions did not reveal obvious repetitive sequences in centromeric regions of genome v3. Hence the under-annotation of TEs in centromeres might also suggest that *A. psidii* centromeres are not enriched in TEs. In future studies, performing centromere analysis with a TE database built on the updated genome and including additional *A. psidii* genomes might resolve this outstanding question.

### **4.4** Signals of extensive genomic degeneration around the *Austropuccinia psidii* PR locus and significant amplification of MFA peptide precursor ORFs

We performed detailed analysis of the *A. psidii* mating-type loci. Our analysis further supports the findings of Ferrarezi et al. (2022) that *A. psidii* has a tetrapolar mating system. Investigation of the full HD locus reveals minimal genomic degeneration while the PR locus exhibit significant degeneration, creating large syntenic gaps. These results are consistent with previous findings for cereal rust fungi (114, 115).

The identification of extensive copies of *mfas* is unexpected, since only 2-3 copies of *mfas* have been identified in other rust fungi (42, 114, 115). These *mfa* duplications and translocations are likely related to TE insertions. Moreover, the high conservation of tandem repeat sequences within *mfa*1/3 across rust fungi, despite substantial evolutionary divergence strongly suggests potential purifying selection on these motifs due to their important function for *STE3.2-2* and *STE3.2-3* pheromone receptor recognition. Additionally, we identified that the PR locus is significantly extended to close to 10 Mb in size which is mostly driven by repeat expansions and likely linked to recombination suppression. At the same time, 5mCpG methylation appears not to be enriched when compared to the rest of the genome and the PR locus in *P. striiformis* f. sp. *tritici* (29).

Mechanisms involved in meiotic recombination suppression in the sex or mating-type regions across organisms include epigenetic modifications, such as DNA cytosine methylation in *Arabidopsis thaliana* (116) and histone H3K9 methylation at the MAT locus in fission yeast, which lacks DNA methylation (117). Given the sequence divergence between the two haplotypes of the PR locus it is likely that sequence heterogeneity shaped by TE insertions inhibits recombination activity at this locus in the absence of DNA methylation.

### 4.5 The pandemic biotype of *Austropuccinia psidii* displays uniform infection dynamics across distantly related plant species

The pandemic lineage of *A. psidii* is linked to the rapid spread and dissemination of the pathogen worldwide (11, 15, 46, 118, 119). This widespread distribution is likely attributable to its polyphagous nature, which allows it to infect a broad range of host species (7). In contrast, the pathogen demonstrates a higher level of host specialization in its centre of origin (10–13). This duality, where the pathogen thrives in its invasive range globally while maintaining specialization in its native range, highlights the complex ecological dynamics at play and underscores the need for further research into its adaptative strategies.

Studies examining the interaction of *A. psidii* with individual host species, have reported vastly different infection time scales on the same host in different laboratories and on different host species across different studies (48, 120, 121). For example, infection dynamics of the susceptible species *S. jambos* ranged from as short as 6 days (48, 121) to 12 days (120) for the completion of the infection cycle. Our research work shows that the pandemic biotype of *A. psidii* displays very consistent infection dynamics across four different host species, despite being separated by 65 mya of evolution (122, 123), when using the same infection conditions. This finding emphasizes the importance of more in depth understanding of infection mechanisms on different host species to develop effective management and biosecurity strategies.

### 4.6 Allele-specific gene expression analysis provides evidence for identifying candidate effectors

While most of the studies investigated the genetic mechanisms of defence of host species against *A. psidii* infection (7, 124–128), only two studies have examined the molecular genetics of the fungus during the infection of susceptible hosts (127, 128), both of which used the early version of the *A. psidii* genome (21). Here, we present the first allele specific expression (ASE) analysis of *A. psidii* over the course of infection in a susceptible host.

The patterns of gene expression clustered into three distinct groups: ungerminated urediniospores, early infection stage (2-4 dpi), and late infection stage (5-6 dpi). These patterns are like those observed in the asexual reproduction of *P. striiformis* f. sp. *tritici* (29, 129). The increase in the number of *A. psidii* DEGs in the late infection stages was also previously reported in the interaction of *A. psidii* with susceptible *E. grandis* and may be attributed to the lack of host defences, which allows the fungus to reproduce more effectively compared to a resistant host (127, 130).

The early upregulation of candidate effectors from 2 to 4 dpi coincides with the latency phase of the disease (before sporulation), while the overexpression of effectors at 5 dpi coincides with the appearance of symptoms. At 6 dpi, several different patterns were observed: some candidate effectors that were previously downregulated were either upregulated or downregulated to a greater extent; while those CEs that were previously upregulated were either downregulated or maintained their upregulation. This timepoint coincides with the first appearance of sporulating lesions, marking a significant transition in the infection cycle. Interestingly, 30 and 25 candidate effectors were only present in hapA or hapB respectively, including two of the top overexpressed genes *in planta*.

Among the overexpressed genes, the majority possess alleles in both haplotypes, with most exhibiting direction-biased expression favouring one allele over the other. In contrast, only a small subset of genes showed consistent expression pattern in both haplotypes. Allele-specific expression may arise from various genomic and epigenomic mechanisms, including somatic copy number alterations (deletions or duplications), gene mutations, epigenetic modifications in promoter regions, and cis-acting regulatory mutations that influence transcriptional outcomes (131). In plants, two categories of ASE genes have been reported: consistent ASE genes, which ASE exhibit bias toward one parental allele across conditions, and inconsistent ASE genes, which ASE display bias toward different parental alleles under varying conditions (132, 133). The latter likely represents adaptive regulation, allowing plants to adjust allelic expression for better response (132, 133). Since the expression difference caused by ASE may result in phenotypic variation (132), future experimental study to verify the CEs might allow better understanding of virulence of *A. psidii*.

## Acknowledgements

Schwessinger and P. A. Tobias were supported by an ARC Linkage grant LP190100093. São Paulo Research Foundation (FAPESP) Grant 2019/13191-5. T. R. Boufleur was supported by FAPESP Grants 2022/11900-1 and 2021/01606-6. We gratefully acknowledge the computational resources provided by the National Computational Infrastructure (NCI) and ANU Merit Allocation Scheme (ANUMAS).

## References

1. Oliver RP. 2024. Diseases caused by fungi, p. 339–428. In Oliver, RP, Huckelhoven, R, Del Ponte, EM, Di Pietro, A (eds.), Agrios’ Plant Pathology, 6th ed. Academic Press.

2. Aime MC, McTaggart AR. 2020. A higher-rank classification for rust fungi, with notes on genera. 10.3114/fuse.2021.07.02.

3. Aime MC, McTaggart AR, Mondo SJ, Duplessis S. 2017. Phylogenetics and Phylogenomics of Rust Fungi. Adv Genet 100:267–307.

4. Ono Y, Buriticá P, Hennen JF. 1992. Delimitation of *Phakopsora, Physopella* and *Cerotelium* and their species on Leguminosae. Mycological Research 96:825– 850.

5. Bonde MR, Nester SE, Berner DK, Frederick RD, Moore WF, Little S. 2008. Comparative Susceptibilities of Legume Species to Infection by Phakopsora pachyrhizi. Plant Dis 92:30–36.

6. Slaminko TL, Miles MR, Frederick RD, Bonde MR, Hartman GL. 2008. New Legume Hosts of Phakopsora pachyrhizi Based on Greenhouse Evaluations. 101094/PDIS-92-5-0767. research-article. The American Phytopathological Society. https://apsjournals.apsnet.org/doi/10.1094/PDIS-92-5-0767. Retrieved 30 January 2025.

7. Soewarto J, Giblin F, Carnegie AJ. 2019. Austropuccinia psidii (myrtle rust) global host list. Version 2. ACT: Australian Network for Plant Conservation, Canberra.

8. Beenken L. 2017. *Austropuccinia*: a new genus name for the myrtle rust *Puccinia psidii* placed within the redefined family Sphaerophragmiaceae (Pucciniales). 1. Phytotaxa 297:53.

9. Ebinghaus M, Gasparotto L, Martins JMT, Santos MDMD, Tessman DJ, Barros-Cordeiro KB, Pinho DB, Dianese JC. 2024. *Austropuccinia licaniae*, first congeneric with the myrtle rust pathogen A. psidii. Mycologia 116:418–430.

10. Graça RN, Ross-Davis AL, Klopfenstein NB, Kim M-S, Peever TL, Cannon PG, Aun CP, Mizubuti ESG, Alfenas AC. 2013. Rust disease of eucalypts, caused by *Puccinia psidii*, did not originate via host jump from guava in Brazil. 24. Mol Ecol 22:6033–6047.

11. Stewart JE, Ross-Davis AL, Graҫa RN, Alfenas AC, Peever TL, Hanna JW, Uchida JY, Hauff RD, Kadooka CY, Kim M -S., Cannon PG, Namba S, Simeto S, Pérez CA, Rayamajhi MB, Lodge DJ, Arguedas M, Medel-Ortiz R, López-Ramirez MA, Tennant P, Glen M, Machado PS, McTaggart AR, Carnegie AJ, Klopfenstein NB. 2018. Genetic diversity of the myrtle rust pathogen ( *Austropuccinia psidii*) in the Americas and Hawaii: Global implications for invasive threat assessments. Forest Pathology 48:e12378.

12. Morales JVP, Boufleur TR, Gonçalves MP, Parisi MCM, Loehrer M, Schaffrath U, Amorim L. 2023. Differential aggressiveness of *Austropuccinia psidii* isolates from guava and rose apple upon cross-inoculation. Plant 10.1111/ppa.13850.

13. Luo Z, Feng J, Bird A, Moeller M, Tam R, Shepherd L, Murphy L, Singh L, Graetz A, Amorim L, Massola Júnior NS, Prodhan MA, Shuey L, Beattie D, Gonzalez AT, Tobias PA, Padovan A, Kimber R, McTaggart A, Kehoe M, Schwessinger B, Boufleur TR. 2025. Plant 109:770–778.

14. Soewarto J, Pérez C, Bartlett M, Somchit C, Ganley B, Sutherland R, Simeto S, Stewart JE, Ibarra Caballero JR, Fraser S, Scott PM, Nadarajan J, Waipara N, Marsh A, Ryan J, Miller E, Smith GR. 2025. New Zealand Myrtaceae are susceptible to a strain from the *Eucalyptus* biotype of *Austropuccinia psidii* present in South America. Biol 27:72.

15. Carnegie AJ, Pegg GS. 2018. Lessons from the Incursion of Myrtle Rust in Australia. Annu Rev Phytopathol 56:457–478.

16. Berthon KA, Fernandez Winzer L, Sandhu K, Cuddy W, Manea A, Carnegie AJ, Leishman MR. 2019. Endangered species face an extra threat: susceptibility to the invasive pathogen *Austropuccinia psidii* (myrtle rust) in Australia. Australasian Plant Pathol 48:385–393.

17. Fensham RJ, Radford-Smith J. 2021. Unprecedented extinction of tree species by fungal disease. Biol Conserv 261:109276.

18. Fensham RJ, Carnegie AJ, Laffineur B, Makinson RO, Pegg GS, Wills J. 2020. Imminent extinction of Australian Myrtaceae by fungal disease. TREE 35:554– 557.

19. Ramos AP, Tavares S, Tavares D, Silva MDC, Loureiro J, Talhinhas P. 2015. Flow cytometry reveals that the rust fungus, Uromyces bidentis (Pucciniales), possesses the largest fungal genome reported--2489 Mbp. Mol Plant Pathol 16:1006–1010.

20. Tavares S, Ramos AP, Pires AS, Azinheira HG, Caldeirinha P, Link T, Abranches R, Silva M do C, Voegele RT, Loureiro J, Talhinhas P. 2014. Genome size analyses of Pucciniales reveal the largest fungal genomes. Front Plant Sci 5.

21. Tobias PA, Schwessinger B, Deng CH, Wu C, Dong C, Sperschneider J, Jones A, Lou Z, Zhang P, Sandhu K, Smith GR, Tibbits J, Chagné D, Park RF. 2021. *Austropuccinia psidii*, causing myrtle rust, has a gigabase-sized genome shaped by transposable elements. G3 11:jkaa015.

22. Wenger AM, Peluso P, Rowell WJ, Chang P-C, Hall RJ, Concepcion GT, Ebler J, Fungtammasan A, Kolesnikov A, Olson ND, Töpfer A, Alonge M, Mahmoud M, Qian Y, Chin C-S, Phillippy AM, Schatz MC, Myers G, DePristo MA, Ruan J, Marschall T, Sedlazeck FJ, Zook JM, Li H, Koren S, Carroll A, Rank DR, Hunkapiller MW. 2019. Accurate circular consensus long-read sequencing improves variant detection and assembly of a human genome. Nat Biotechnol 37:1155–1162.

23. Li H, Durbin R. 2024. Genome assembly in the telomere-to-telomere era. Nat Rev Genet 25:658–670.

24. Duan H, Jones AW, Hewitt T, Mackenzie A, Hu Y, Sharp A, Lewis D, Mago R, Upadhyaya NM, Rathjen JP, Stone EA, Schwessinger B, Figueroa M, Dodds PN, Periyannan S, Sperschneider J. 2022. Physical separation of haplotypes in dikaryons allows benchmarking of phasing accuracy in Nanopore and HiFi assemblies with Hi-C data. Genome Biology 23:84.

25. Henningsen EC, Hewitt T, Dugyala S, Nazareno ES, Gilbert E, Li F, Kianian SF, Steffenson BJ, Dodds PN, Sperschneider J, Figueroa M. 2022. A chromosome-level, fully phased genome assembly of the oat crown rust fungus *Puccinia* sp. *avenae*: a resource to enable comparative genomics in the cereal rusts. G3 12:jkac149.

26. Schwessinger B, Jones A, Albekaa M, Hu Y, Mackenzie A, Tam R, Nagar R, Milgate A, Rathjen JP, Periyannan S. 2022. A chromosome scale assembly of a australian *Puccinia striiformis* f. sp. *tritici* isolate of the PstS1 lineage. MPMI 35:293–296.

27. Li C, Qiao L, Lu Y, Xing G, Wang X, Zhang G, Qian H, Shen Y, Zhang Y, Yao W, Cheng K, Ma Z, Liu N, Wang D, Zheng W. 2023. Gapless genome assembly of *Puccinia triticina* provides insights into chromosome evolution in Pucciniales. 11:e02828-22.

28. Liang J, Li Y, Dodds PN, Figueroa M, Sperschneider J, Han S, Tsui CKM, Zhang K, Li L, Ma Z, Cai L. 2023. Haplotype-phased and chromosome-level genome assembly of *Puccinia polysora*, a giga-scale fungal pathogen causing southern corn rust.

29. Tam R, Möller M, Luo R, Luo Z, Jones A, Periyannan S, Rathjen JP, Schwessinger B. 2024. Long-read genomics reveal extensive nuclear-specific evolution and allele-specific expression in a dikaryotic fungus. bioRxiv 10.1101/2024.12.11.628074.

30. Sperschneider J, Chen J, Anderson C, Morin E, Zhang X, Lewis D, Henningsen E, Grigoriev IV, Rathjen JP, Jones DA, Duplessis S, Dodds PN. 2025. A chromosome-scale genome assembly of the flax rust fungus reveals the two unusually large effector proteins, AvrM3 and AvrN. bioRxiv 10.1101/2025.04.28.651126.

31. Fouché S, Oggenfuss U, Chanclud E, Croll D. 2022. A devil’s bargain with transposable elements in plant pathogens. Trends 38:222–230.

32. Bourgeois Y, Boissinot S. 2019. On the Population Dynamics of Junk: A on the of Genes 10:419.

33. Mita P, Boeke JD. 2016. How retrotransposons shape genome regulation. Curr Opin Genet Dev 37:90–100.

34. Selker EU. 1990. Premeiotic instability of repeated sequences in *Neurospora crassa*. Annu Rev Genet 24:579–613.

35. Selker EU. 2002. Repeat-induced gene silencing in fungi. Adv Genet 46:439– 450.

36. Buchon N, Vaury C. 2006. RNAi: a defensive RNA-silencing against viruses and transposable elements. Heredity 96:195–202.

37. Zhou W, Liang G, Molloy PL, Jones PA. 2020. DNA methylation enables transposable element-driven genome expansion.

38. Luo Z, McTaggart A, Schwessinger B. 2024. Genome biology and evolution of mating-type loci in four cereal rust fungi. 20:e1011207.

39. Turgeon BG. 1998. Application of mating type gene technology to problems in fungal biology. Annu Rev Phytopathol 36:115–137.

40. Coelho MA, Bakkeren G, Sun S, Hood ME, Giraud T. 2017. Fungal sex: the Basidiomycota. Microbiol Spectr 5.

41. Hsueh Y-P, Heitman J. 2008. Orchestration of sexual reproduction and virulence by the fungal mating-type locus.

42. Cuomo CA, Bakkeren G, Khalil HB, Panwar V, Joly D, Linning R, Sakthikumar S, Song X, Adiconis X, Fan L, Goldberg JM, Levin JZ, Young S, Zeng Q, Anikster Y, Bruce M, Wang M, Yin C, McCallum B, Szabo LJ, Hulbert S, Chen X, Fellers JP. 2017. Comparative analysis highlights variable genome content of wheat rusts and divergence of the mating loci.

43. Edwards RJ, Dong C, Park RF, Tobias PA. 2022. A phased chromosome-level genome and full mitochondrial sequence for the dikaryotic myrtle rust pathogen, Austropuccinia psidii. bioRxiv 10.1101/2022.04.22.489119.

44. Ferrarezi JA, McTaggart AR, Tobias PA, Hayashibara CAA, Degnan RM, Shuey LS, Franceschini LM, Lopes MS, Quecine MC. 2022. *Austropuccinia psidii* uses tetrapolar mating and produces meiotic spores in older infections on *Eucalyptus grandis*. Fungal 160:103692.

45. Boufleur TR, Morales JVP, Martins TV, Gonçalves MP, Júnior NSM, Amorim L. 2023. A diagnostic guide for myrtle rust. Plant Health 24:242–251.

46. Ruffner B, Beenken L, Kupper Q, Mittelstrass J, Schuler P, Stewart JE, Caballero JRI, Winiger R, Prospero S. 2024. First report of *Austropuccinia psidii* on *Syzygium buxifolium* grown as indoor bonsai in Europe. New Disease Reports 50:e70011.

47. Yong WTL, Ades PK, Tibbits JFG, Bossinger G, Runa FA, Sandhu KS, Taylor PWJ. 2019. Disease cycle of *Austropuccinia psidii* on *Eucalyptus globulus* and *Eucalyptus obliqua* leaves of different rust response phenotypes. 3. Plant Pathol 68:547–556.

48. Beresford RM, Shuey LS, Pegg GS. 2020. Symptom development and latent period of *Austropuccinia psidii* (myrtle rust) in relation to host species, temperature, and ontogenic resistance. 3. Plant 69:484–494.

49. Smith GR, Ganley BJ, Chagné D, Nadarajan J, Pathirana RN, Ryan J, Arnst EA, Sutherland R, Soewarto J, Houliston G, Marsh AT, Koot E, Carnegie AJ, Menzies T, Lee DJ, Shuey LS, Pegg GS. 2020. Resistance of New Zealand Provenance *Leptospermum scoparium, Kunzea robusta, Kunzea linearis* , and *Metrosideros excelsa* to *Austropuccinia psidii*. Plant Dis 104:1771–1780.

50. Martino AM, Park RF, Tobias PA. 2024. Threatened and priority listed *Melaleuca* species from Western Australia display high susceptibility to *Austropuccinia psidii* in controlled inoculations. Australasian Plant Pathol 53:253–260.

51. Morin L, Aveyard R, Lidbetter R, Wilson PG. 2012. Investigating the host-range of the rust fungus *Puccinia psidii* sensu lato across tribes of the family Myrtaceae present in Australia. PLoS ONE 7:e35434.

52. Chen SH, Yap J-YS, Viler V, Stehn C, Sandhu KS, Percival J, Pegg GS, Menzies T, Jones A, Guo K, Giblin FR, Cohen J, Edwards RJ, Rossetto M, Bragg JG. 2024. Genomics and resistance assays inform the management of two tree species being devastated by the invasive myrtle rust pathogen 10.1101/2024.10.30.612564.

53. Schwessinger B. 2016. High quality DNA from Fungi for long read sequencing e.g. PacBio. protocols.io. https://www.protocols.io/view/High-quality-DNA-from-Fungi-for-long-read-sequenci-ewtbfen. Retrieved 1 September 2022.

54. Baid G, Cook DE, Shafin K, Yun T, Llinares-López F, Berthet Q, Belyaeva A, Töpfer A, Wenger AM, Rowell WJ, Yang H, Kolesnikov A, Ammar W, Vert J-P, Vaswani A, McLean CY, Nattestad M, Chang P-C, Carroll A. 2023. DeepConsensus improves the accuracy of sequences with a gap-aware sequence transformer. 2. Nat Biotechnol 41:232–238.

55. De Coster W, Rademakers R. 2023. NanoPack2: population-scale evaluation of long-read sequencing data. Bioinformatics 39:btad311.

56. Inglis PW, Pappas M de CR, Resende LV, Grattapaglia D. 2018. Fast and inexpensive protocols for consistent extraction of high quality DNA and RNA from challenging plant and fungal samples for high-throughput SNP genotyping and sequencing applications. ONE 13:e0206085.

57. Chang S, Puryear J, Cairney J. 1993. A simple and efficient method for isolating RNA from pine trees. Plant Mol Biol Rep 11:113–116.

58. Cheng H, Concepcion GT, Feng X, Zhang H, Li H. 2021. Haplotype-resolved de novo assembly using phased assembly graphs with hifiasm. Nat Methods 18:170–175.

59. Cheng H, Jarvis ED, Fedrigo O, Koepfli K-P, Urban L, Gemmell NJ, Li H. 2022. Haplotype-resolved assembly of diploid genomes without parental data. Nat Biotechnol 40:1332–1335.

60. Li H. 2021. New strategies to improve minimap2 alignment accuracy. Bioinformatics 37:4572–4574.

61. Danecek P, Bonfield JK, Liddle J, Marshall J, Ohan V, Pollard MO, Whitwham A, Keane T, McCarthy SA, Davies RM, Li H. 2021. Twelve years of SAMtools and BCFtools. GigaScience 10:giab008.

62. Durand NC, Shamim MS, Machol I, Rao SSP, Huntley MH, Lander ES, Aiden EL. 2016. Juicer provides a one-click system for analyzing loop-resolution Hi-C experiments. Cell Syst 3:95–98.

63. Durand NC, Robinson JT, Shamim MS, Machol I, Mesirov JP, Lander ES, Aiden EL. 2016. Juicebox Provides a Visualization System for Hi-C Contact Maps with Unlimited Zoom. Cell Syst 3:99–101.

64. Simão FA, Waterhouse RM, Ioannidis P, Kriventseva EV, Zdobnov EM. 2015. BUSCO: assessing genome assembly and annotation completeness with single-copy orthologs. Bioinformatics 31:3210–3212.

65. Field MA, Rosen BD, Dudchenko O, Chan EKF, Minoche AE, Edwards RJ, Barton K, Lyons RJ, Tuipulotu DE, Hayes VM, D. Omer A, Colaric Z, Keilwagen J, Skvortsova K, Bogdanovic O, Smith MA, Aiden EL, Smith TPL, Zammit RA, Ballard JWO. 2020. Canfam_GSD: De novo chromosome-length genome assembly of the German Shepherd Dog (*Canis lupus familiaris*) using a combination of long reads, optical mapping, and Hi-C. GigaScience 9:giaa027.

66. Emms DM, Kelly S. 2015. OrthoFinder: solving fundamental biases in whole genome comparisons dramatically improves orthogroup inference accuracy. 1. Genome Biology 16:157.

67. Wu T, Hu E, Xu S, Chen M, Guo P, Dai Z, Feng T, Zhou L, Tang W, Zhan L, Fu X, Liu S, Bo X, Yu G. 2021. clusterProfiler 4.0: A universal enrichment tool for interpreting omics data. The Innovation 2:100141.

68. Teufel F, Almagro Armenteros JJ, Johansen AR, Gíslason MH, Pihl SI, Tsirigos KD, Winther O, Brunak S, von Heijne G, Nielsen H. 2022. SignalP 6.0 predicts all five types of signal peptides using protein language models. Nat Biotechnol 40:1023–1025.

69. Hallgren J, Tsirigos KD, Pedersen MD, Armenteros JJA, Marcatili P, Nielsen H, Krogh A, Winther O. 2022. DeepTMHMM predicts alpha and beta transmembrane proteins using deep neural networks. bioRxiv 10.1101/2022.04.08.487609.

70. Gíslason MH, Nielsen H, Almagro Armenteros JJ, Johansen AR. 2021. Prediction of GPI-anchored proteins with pointer neural networks.

71. Horton P, Park K-J, Obayashi T, Fujita N, Harada H, Adams-Collier CJ, Nakai K. 2007. WoLF PSORT: protein localization predictor. Nucleic Acids Res 35:W585–587.

72. Altschul SF, Gish W, Miller W, Myers EW, Lipman DJ. 1990. Basic local alignment search tool. J Mol Biol 215:403–410.

73. Johnson M, Zaretskaya I, Raytselis Y, Merezhuk Y, McGinnis S, Madden TL. 2008. NCBI BLAST: a better web interface. Nucleic Acids Res 36:W5–9.

74. Kolde R. 2019. pheatmap: Pretty Heatmaps (1.0.12).

75. Li F, Upadhyaya NM, Sperschneider J, Matny O, Nguyen-Phuc H, Mago R, Raley C, Miller ME, Silverstein KAT, Henningsen E, Hirsch CD, Visser B, Pretorius ZA, Steffenson BJ, Schwessinger B, Dodds PN, Figueroa M. 2019. Emergence of the Ug99 lineage of the wheat stem rust pathogen through somatic hybridisation. Nat Commun 10:5068.

76. Hunter JD. 2007. Matplotlib: A 2D Graphics Environment.

77. Lawrence M, Huber W, Pagès H, Aboyoun P, Carlson M, Gentleman R, Morgan MT, Carey VJ. 2013. Software for computing and annotating genomic ranges. PLoS Comput Biol 9:e1003118.

78. Gel B, Serra E. 2017. karyoploteR: an R/Bioconductor package to plot customizable genomes displaying arbitrary data. Bioinformatics 33:3088–3090.

79. Upadhyay UD, Dworkin SL, Weitz TA, Foster DG. 2014. Development and validation of a reproductive autonomy scale. Stud Fam Plann 45:19–41.

80. Virtanen P, Gommers R, Oliphant TE, Haberland M, Reddy T, Cournapeau D, Burovski E, Peterson P, Weckesser W, Bright J, van der Walt SJ, Brett M, Wilson J, Millman KJ, Mayorov N, Nelson ARJ, Jones E, Kern R, Larson E, Carey CJ, Polat İ, Feng Y, Moore EW, VanderPlas J, Laxalde D, Perktold J, Cimrman R, Henriksen I, Quintero EA, Harris CR, Archibald AM, Ribeiro AH, Pedregosa F, van Mulbregt P, SciPy 1.0 Contributors. 2020. SciPy 1.0: fundamental algorithms for scientific computing in Python. Nat Methods 17:261–272.

81. Quinlan AR, Hall IM. 2010. BEDTools: a flexible suite of utilities for comparing genomic features. Bioinformatics 26:841–842.

82. Dale RK, Pedersen BS, Quinlan AR. 2011. Pybedtools: a flexible Python library for manipulating genomic datasets and annotations. Bioinformatics 27:3423– 3424.

83. Waskom ML. 2021. seaborn: statistical data visualization.

84. Sievers F, Higgins DG. 2018. Clustal Omega for making accurate alignments of many protein sequences. Protein Sci 27:135–145.

85. Cao T, Li Q, Huang Y, Li A. 2023. plotnineSeqSuite: a Python package for visualizing sequence data using ggplot2 style. BMC 24:585.

86. Lechner M, Findeiß S, Steiner L, Marz M, Stadler PF, Prohaska SJ. 2011. Proteinortho: Detection of (Co-)orthologs in large-scale analysis. BMC 12:124.

87. Edgar RC. 2004. MUSCLE: multiple sequence alignment with high accuracy and high throughput. Nucleic Acids Res 32:1792–1797.

88. Yang Z. 2007. PAML 4: phylogenetic analysis by maximum likelihood. Mol Biol Evol 24:1586–1591.

89. Kim D, Paggi JM, Park C, Bennett C, Salzberg SL. 2019. Graph-based genome alignment and genotyping with HISAT2 and HISAT-genotype. Nat Biotechnol 37:907–915.

90. Patro R, Duggal G, Love MI, Irizarry RA, Kingsford C. 2017. Salmon provides fast and bias-aware quantification of transcript expression. Nat Methods 14:417– 419.

91. R Core Team. 2024. A Language and Environment for Statistical Computing. R Foundation for Statistical Computing. R. R Foundation for Statistical Computing, Vienna, Austria.

92. Soneson C, Love MI, Robinson MD. 2016. Differential analyses for RNA-seq: transcript-level estimates improve gene-level inferences. 4:1521. F1000Research 10.12688/f1000research.7563.1.

93. Robinson MD, McCarthy DJ, Smyth GK. 2010. edgeR: a Bioconductor package for differential expression analysis of digital gene expression data. Bioinformatics 26:139–140.

94. Ritchie ME, Phipson B, Wu D, Hu Y, Law CW, Shi W, Smyth GK. 2015. limma powers differential expression analyses for RNA-sequencing and microarray studies. Nucleic Acids 43:e47.

95. Berger BA, Kriebel R, Spalink D, Sytsma KJ. 2016. Divergence times, historical biogeography, and shifts in speciation rates of Myrtales. Mol Phylogenet Evol 95:116–136.

96. Boehm E, Bushnell W. 1992. An ultrastructural pachytene karyotype for *Melampsora lini*. PHYTOPATHOLOGY-NEW YORK AND BALTIMORE THEN ST PAUL 82:1212–1212.

97. Boehm E, Wenstrom J, Mclaughlin D, Szabo L, Roelfs A, Bushnell W. 1992. An ultrastructural pachytene karyotype for *Puccinia graminis* f. sp. *tritici*. Canadian Journal of Botany 70:401–403.

98. Aime MC, Bell CD, Wilson AW. 2018. Deconstructing the evolutionary complexity between rust fungi (*Pucciniales*) and their plant hosts. Studies in Mycology 89:143–152.

99. Goddard MV. 1976. Cytological studies of *Puccinia striiformis* (yellow rust of wheat).

100. Valkoun J, Bartoš P. 1974. Somatic chromosome number in *Puccinia recondita*.

101. Savile DB. 1939. Nuclear structure and behavior in species of the Uredinales. American 585–609.

102. Moynahan ME, Jasin M. 2010. Mitotic homologous recombination maintains genomic stability and suppresses tumorigenesis. Nat Rev Mol Cell Biol 11:196– 207.

103. Ip K, Yadin R, George KW. 2020. High-Throughput DNA Assembly Using Yeast Homologous Recombination, p. 79–89. *In* Chandran, S, George, KW (eds.), DNA Cloning and Assembly: Methods and Protocols. Springer US, New York, NY.

104. Xu Y, Tian L, Tan J, Huang W, Li J, O’Neil N, Hirst M, Hieter P, Zhang Y, Li X. 2025. Distribution of haploid chromosomes into separate nuclei in two pathogenic fungi. Science 388:784–788.

105. Rustchenko E. 2007. Chromosome instability in *Candida albicans*. FEMS Yeast Research 7:2–11.

106. Rancati G, Pavelka N, Fleharty B, Noll A, Trimble R, Walton K, Perera A, Staehling-Hampton K, Seidel CW, Li R. 2008. Aneuploidy underlies rapid adaptive evolution of yeast cells deprived of a conserved cytokinesis motor. Cell 135:879–893.

107. Sah SK, Hayes JJ, Rustchenko E. 2021. The role of aneuploidy in the emergence of echinocandin resistance in human fungal pathogen *Candida albicans*. 17:e1009564.

108. Bird AP. 1980. DNA methylation and the frequency of CpG in animal DNA. Nucleic Acids Res 8:1499–1504.

109. Bird AP, Taggart MH. 1980. Variable patterns of total DNA and rDNA methylation in animals. Nucleic Acids Res 8:1485–1497.

110. Simmonds P, Xia W, Baillie JK, McKinnon K. 2013. Modelling mutational and selection pressures on dinucleotides in eukaryotic phyla--selection against CpG and UpA in cytoplasmically expressed RNA and in RNA viruses. BMC 14:610.

111. Ying H, Hayward DC, Klimovich A, Bosch TCG, Baldassarre L, Neeman T, Forêt S, Huttley G, Reitzel AM, Fraune S, Ball EE, Miller DJ. 2022. The role of DNA methylation in genome defense in Cnidaria and other invertebrates. Mol Biol Evol 39:msac018.

112. Sperschneider J, Jones AW, Nasim J, Xu B, Jacques S, Zhong C, Upadhyaya NM, Mago R, Hu Y, Figueroa M, Singh KB, Stone EA, Schwessinger B, Wang M-B, Taylor JM, Dodds PN. 2021. The stem rust fungus *Puccinia graminis* f. sp. *tritici* induces centromeric small RNAs during late infection that are associated with genome-wide DNA methylation. BMC Biol 19:203.

113. Hernández-Saavedra D, Strakovsky RS, Ostrosky-Wegman P, Pan Y-X. 2017. Epigenetic regulation of centromere chromatin stability by dietary and environmental factors.

114. Henningsen EC, Lewis D, Nazareno ES, Mangelson H, Sanchez M, Langford K, Huang Y-F, Steffenson BJ, Boesen B, Kianian SF, Liachko I, Stone E, Dodds PN, Sperschneider J, Figueroa M. 2024. A high-resolution haplotype collection uncovers somatic hybridization, recombination and intercontinental movement in oat crown rust. Genetics 20:e1011493.

115. Luo Z, McTaggart A, Schwessinger B. 2023. Genome biology and evolution of mating type loci in four cereal rust fungi. bioRxiv 10.1101/2023.03.02.530769.

116. Ge T, Gui X, Xu J-X, Xia W, Wang C-H, Yang W, Huang K, Walsh C, Umen JG, Walter J, Du Y-R, Chen H, Shao Z, Xu G-L. 2024. DNA cytosine methylation suppresses meiotic recombination at the sex-determining region. Sci Adv 10:eadr2345.

117. Oh J, Yeom S, Park J, Lee J-S. 2022. The regional sequestration of heterochromatin structural proteins is critical to form and maintain silent chromatin. Epigenetics Chromatin 15:5.

118. Granados GM, McTaggart AR, Barnes I, Rodas CA, Roux J, Wingfield MJ. 2017. The pandemic biotype of *Austropuccinia psidii* discovered in South America. Australasian Plant Pathol 46:267–275.

119. du Plessis E, Granados GM, Barnes I, Ho WH, Alexander BJR, Roux J, McTaggart AR. 2019. The pandemic strain of *Austropuccinia psidii* causes myrtle rust in New Zealand and Singapore. 3. Australasian Plant Pathol 48:253–256.

120. Glen M, Alfenas AC, Zauza EAV, Wingfield MJ, Mohammed C. 2007. *Puccinia psidii* : a threat to the Australian environment and economy – a review. 1. Austral Plant Pathol 36:1.

121. Degnan RM, Shuey LS, Radford-Smith J, Gardiner DM, Carroll BJ, Mitter N, McTaggart AR, Sawyer A. 2023. Double-stranded RNA prevents and cures infection by rust fungi. Commun Biol 6:1–10.

122. Grattapaglia D, Vaillancourt RE, Shepherd M, Thumma BR, Foley W, Külheim C, Potts BM, Myburg AA. 2012. Progress in Myrtaceae genetics and genomics: *Eucalyptus* as the pivotal genus. Tree Genetics & Genomes 8:463–508.

123. Biffin E, Lucas EJ, Craven LA, Ribeiro da Costa I, Harrington MG, Crisp MD. 2010. Evolution of exceptional species richness among lineages of fleshy-fruited Myrtaceae. Ann Bot 106:79–93.

124. Hsieh J-F, Chuah A, Patel HR, Sandhu KS, Foley WJ, Külheim C. 2018. Transcriptome profiling of *Melaleuca quinquenervia* challenged by myrtle rust reveals differences in defense responses among resistant individuals. Phytopathology® 108:495–509.

125. Tobias PA, Guest DI, Külheim C, Park RF. 2018. *De Novo* transcriptome study identifies candidate genes involved in resistance to *Austropuccinia psidii* (myrtle Rust) in *Syzygium luehmannii* (riberry). 108:627–640.

126. Santos SA, Vidigal PMP, Guimarães LMS, Mafia RG, Templeton MD, Alfenas AC. 2020. Transcriptome analysis of *Eucalyptus grandis* genotypes reveals constitutive overexpression of genes related to rust (*Austropuccinia psidii*) resistance. Plant Mol Biol 104:339–357.

127. Swanepoel S, Oates CN, Shuey LS, Pegg GS, Naidoo S. 2021. Transcriptome analysis of *Eucalyptus grandis* implicates brassinosteroid signaling in defense against myrtle Rust (*Austropuccinia psidii*).

128. Frampton RA, Shuey LS, David CC, Pringle GM, Kalamorz F, Pegg GS, Chagné D, Smith GR. 2024. Analysis of plant and fungal transcripts from resistant and susceptible phenotypes of *Leptospermum scoparium* challenged by *Austropuccinia psidii*. 114:2121–2130.

129. Schwessinger B, Sperschneider J, Cuddy WS, Garnica DP, Miller ME, Taylor JM, Dodds PN, Figueroa M, Park RF, Rathjen JP. 2018. A near-complete haplotype-phased genome of the dikaryotic wheat stripe rust fungus *Puccinia striiformis f. sp. tritici* reveals high interhaplotype diversity. mBio 9:e02275–17.

130. Swanepoel S, Visser EA, Shuey L, Naidoo S. 2023. The *in planta* gene expression of *Austropuccinia psidii* in resistant and susceptible *Eucalyptus grandis*. PHYTO-07-22-0257-R.

131. Sen A, Prager BC, Zhong C, Park D, Zhu Z, Gimple RC, Wu Q, Bernatchez JA, Beck S, Clark AE, Siqueira-Neto JL, Rich JN, McVicker G. 2022. Leveraging allele-specific expression for therapeutic response gene discovery in glioblastoma. Cancer Res 82:377–390.

132. Shao L, Xing F, Xu C, Zhang Q, Che J, Wang X, Song J, Li X, Xiao J, Chen L-L, Ouyang Y, Zhang Q. 2019. Patterns of genome-wide allele-specific expression in hybrid rice and the implications on the genetic basis of heterosis.

133. Zhan W, Cui L, Yang S, Zhang K, Zhang Y, Yang J. 2024. Natural variations of heterosis-related allele-specific expression genes in promoter regions lead to allele-specific expression in maize. BMC 25:476.

134. Rueden CT, Schindelin J, Hiner MC, DeZonia BE, Walter AE, Arena ET, Eliceiri KW. 2017. ImageJ2: ImageJ for the next generation of scientific image data. BMC Bioinformatics 18:529.

135. Davey NE, Edwards RJ, Shields DC. 2007. The SLiMDisc server: short, linear motif discovery in proteins. Nucleic Acids Res 35:W455–W459.

136. Almeida JR de, Pachón DMR, Franceschini LM, Santos IB dos, Ferrarezi JA, Andrade PAM de, Monteiro-Vitorello CB, Labate CA, Quecine MC. 2021. Revealing the high variability on nonconserved core and mobile elements of *Austropuccinia psidii* and other rust mitochondrial genomes. PLOS ONE 16:e0248054.

137. Tillich M, Lehwark P, Pellizzer T, Ulbricht-Jones ES, Fischer A, Bock R, Greiner S. 2017. GeSeq – versatile and accurate annotation of organelle genomes. Nucleic Acids Research 45:W6–W11.

